# Pervasive Under-Dominance in Gene Expression Underlying Emergent Growth Trajectories in *Arabidopsis thaliana* Hybrids

**DOI:** 10.1101/2022.03.03.482808

**Authors:** Wei Yuan, Fiona Beitel, Thanvi Srikant, Ilja Bezrukov, Sabine Schäfer, Robin Kraft, Detlef Weigel

**Affiliations:** Department of Molecular Biology, Max Planck Institute for Biology Tübingen, 72076 Tübingen, Germany

## Abstract

Complex traits, such as growth and fitness, are typically controlled by a very large number of variants, which can interact in both additive and non-additive fashion. In an attempt to gauge the relative importance of both types of genetic interactions, we have turned to hybrids, which provide a facile means for creating many novel allele combinations. We focused on the interaction between alleles of the same locus and performed a transcriptomic study involving 141 random crosses between different accessions of the plant model species *Arabidopsis thaliana*. Additivity is rare, consistently observed for only about 300 genes enriched for roles in stress response and cell death. Regulatory rare-allele burden affects the expression level of these genes but does not correlate with F_1_ rosette size. Non-additive gene expression in F_1_ hybrids is much more common, with the vast majority of genes (over 90%) being expressed below parental average. Unlike in the additive genes, regulatory rare-allele burden in the non-additive gene set is strongly correlated with F_1_ rosette size, even though it only mildly covary with the expression level of these genes. Our study underscores under-dominance as the predominant gene action associated with emergence of rosette growth trajectories in the *A. thaliana* hybrid model. Our work lays the foundation for understanding molecular mechanisms and evolutionary forces that lead to dominance complementation of rare regulatory alleles.

## Introduction

When expression and inheritance of a trait is under control of many genes, it is considered a quantitative or complex trait, with growth- and fitness-related traits being almost always complex traits (Braendle et al. 2011; Falconer and Mackay 1996). The complexity of these traits comes not only from the large number of underlying genes/loci, but also from the multitude of potential allelic interactions within and between the genes involved. The best-understood type of allelic interaction is the additive effect, in which different alleles contribute to a trait co-dominantly, and the offspring have an intermediate trait value that is close to the average of the two parental alleles. By definition, non-additive genetic effects are any deviations from this additive scenario, with two common examples being dominance and epistasis (Falconer and Mackay 1996). Due to limitations in technical and statistical frameworks, non-additive effects are much less studied and understood than additive effects (Varona et al. 2018). Nevertheless, there is evidence that non-additive effects can be pervasive and contribute substantially to what had for some time been perceived as “missing-heritability” (Mackay et al. 2009; Zuk et al. 2012; Hemani et al. 2013). Additive and non-additive genetic effects are best revealed in heterozygotes, in which contributions of different alleles to a phenotype of interest can be easily partitioned.

Causal genetic variants often exert their functional effects by modulating gene expression. Measuring gene expression differences and associating the variation with complex traits provides information regarding biological functions and processes causing natural phenotypic variation (Albert and Kruglyak 2015). Over the last decade, statistical frameworks such as Transcriptome-wide association (TWA) and expression QTL (eQTL) have rapidly matured, providing insights into molecular functions of complex traits (Albert and Kruglyak 2015; Wainberg et al. 2019). The highly quantitative nature of transcriptomic, especially RNA-seq data provides an excellent opportunity for tracking additive versus non-additive gene actions. By comparing the expression level of a gene in F_1_ heterozygotes to that of the parental average, one can easily calculate the degree of its non-additivity (Jeffrey Chen 2013).

Above-ground biomass accumulation is a fitness-related trait that has important bearing on both local adaptation of wild plants (Vogel et al. 2019) and performance of agricultural species (Batts et al. 1997; Elings et al. 1997). In the inbreeding plant *Arabidopsis thaliana*, rosette-size, a close proxy for above-ground biomass, is not only a primary indicator of growth and general performance (Julkowska et al. 2016; González et al. 2020), but also both highly variable (González et al. 2020; Wieters et al. 2021) and strongly associated with fitness (Korves et al. 2007; Wieters et al. 2021). The range of variation in this trait can be substantially increased by including F_1_ hybrids. While rare combinations are smaller than any naturally occurring accession (Bomblies and Weigel 2007; Chae et al. 2014), most F_1_ hybrids have larger rosettes (Seymour et al. 2016; Yang et al. 2017; Oakley et al. 2019), an emergent positive phenotype that in outbreeding crops is usually termed heterosis. F_1_ hybrids are a particularly interesting systems to study, as alleles that are naturally segregated into different genomes are brought into contact with each other, leading to numerous novel genetic interactions (Landry et al. 2007; Rieseberg et al. 2007). It has long been postulated that these novel genetic interactions, both additive and non-additive, may contribute to hybrid performance (Birchler et al. 2003; Rieseberg et al. 2007; Bell et al. 2013).

We are interested in understanding how common additive and non-additive gene action is, and how it relates, if at all, to growth phenotypes in F_1_ hybrids. Specifically, we would like to learn at the species level (i) whether additive and non-additive gene actions occur at similar frequency, (ii) whether the gene actions are mostly specific to parental combinations, or if certain genes and pathways particularly frequently exhibit one of the effects, (iii) and whether emergent phenotypes in F_1_ hybrids are more likely to result from additive or non-additive gene actions. *Arabidopsis thaliana* provides a powerful system to address these questions, due to the wealth of genomic resources and large collection of natural accessions (1001 Genomes Consortium). Previous studies of intra-specific *A. thalian* hybrids have provided insights into mechanisms affecting hybrid performance, such as the mitigation of defense-growth tradeoffs in superior hybrids (Miller et al. 2015), and on the flip side, greatly compromised growth in hybrids due to incompatible allelic interaction and excessive activation of defense (Chae et al. 2014).

We designed a study in *A. thaliana* that surveyed not only a broad range of the species’ genetic diversity, but also allowed for the detection of interactions between an exceptionally large number of alleles. We find non-additivity in gene expression in F_1_ hybrids to be common, with non-additive genes being much more commonly expressed below the parental average (mid-parent value, MPV) than above it. Expression close to the MPV in turn is rare, with a substantial fraction of such genes having a role in biotic defense pathways, suggesting that defense is particularly well buffered.

## Results

### Non-additive gene action is more abundant in F_1_ hybrids

For our work, we drew on resources from the 1001 Genomes Project for this species (1001 Genomes Consortium), crossing resequenced, naturally inbred accessions to generate a panel of F_1_ hybrids. To broadly survey possible genetic interactions, and to evaluate whether consistent patterns of additive and non-additive gene expression exist, we carried out an RNA-seq experiment for which 101 parent-F_1_ trios, *i*.*e*., each F_1_ hybrid and their inbred parents, were planted (Fig. 1A, Data S1, Methods). Of all genotypes, 82 F_1_s and 124 inbred parents were included in the final analyses. Whole-rosette sizes, a good proxy for biomass (Fig. S1, Methods), were measured in inbreds and F_1_s.

**Figure 1.**
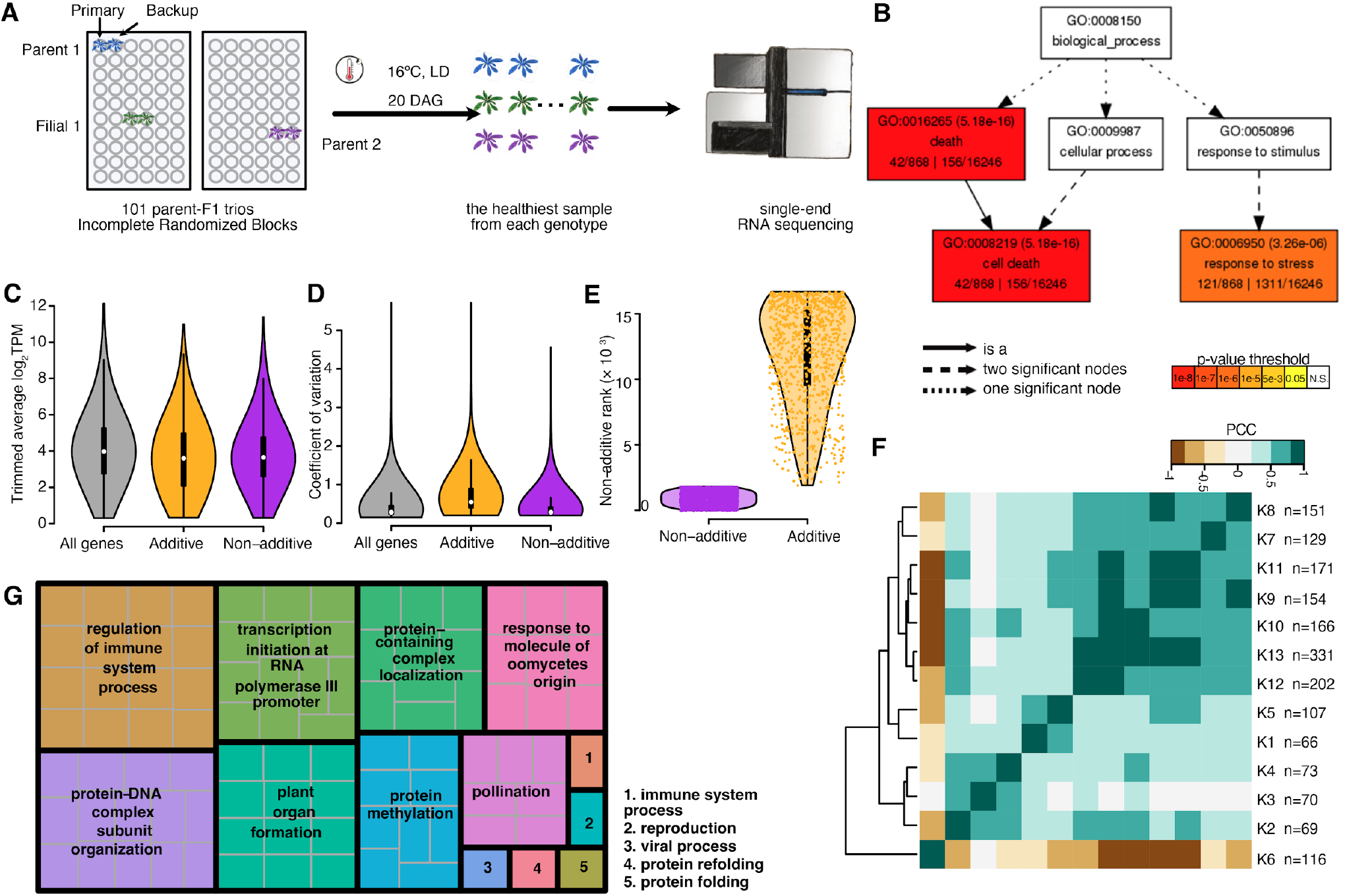
Summary of additive and non-additive genes. **A**. Experimental set-up. Note that not all trios were completely sequenced and analyzed. **B**. GO-term (biological process) enrichment of additive genes. **C-D**. Both additive and non-additive genes showed average transcript abundance (C) and coefficient of variation (D) profiles comparable to those of all genes in the background. **E**. Additive genes ranked low by their non-additive score. **F**. Correlation among all non-additive gene clusters. Pearson correlation coefficients were calculated using cluster average. **G**. Revigo summary of biological processes enriched in non-additive genes.

The expression of many genes changes in F_1_ hybrids relative to the parents (Jeffrey Chen 2013). Under an additive model, gene expression in F_1_ hybrids is close to the parental average. We asked whether there are genes that are almost always additively, or non-additively expressed in F_1_s across all trios. We identified close to 900 genes that were consistently expressed in an additive fashion (Fig. S2, Methods), and 1,805 genes (Data S2) that were consistently expressed non-additively (Fig. S3, Methods) in all trios. Both the additive and the non-additive genes had distribution profiles of transcript abundances and coefficients of variation that were similar to background genes (Fig.1C-D), *i*.*e*., including both high- and low-abundance transcripts as well as ones that varied little across samples or ones that varied substantially. This indicates that expression level and variance did not greatly bias our ability to discover specific gene expression patterns. Parenthetically, the great majority of the additive genes had very low ranks on their non-additive score (Fig. 1E), confirming that our gene calling algorithm successfully selected for contrasting gene actions. Within our non-additive genes, only 150 (∼8%) of these were consistently expressed above MPV in the F_1_s. For the great majority of genes, low expression level was dominant, such that most genes were expressed in the F_1_s at a level between the lower parent and the MPV.

Gene Ontology (GO) term analysis revealed that the additive genes are strongly enriched in cell death- and stress response-related processes (Fig. 1B). Because expression level close to parental average would be indicative of canalization, our result suggests that the cell death- and stress-response pathways are systematically buffered in F_1_s. In comparison, non-additive genes are featured in much more diverse biological processes (Fig. 1G), including regulation of immune system process, ribosomal RNA transcription, plant organ formation, and others.

Non-additive gene expression is thus pervasive in F_1_ hybrids, with the overwhelming majority of genes expressed below the parental average. Many biological processes are affected in the F_1_s by non-additive expression, as the GO enrichment suggested.

### Non-additive genes covary with size

We reduced the dimensionality of our non-additive gene set by grouping genes with a similar behavior across samples via k-means clustering (k=13). We examined how well the behavior of different clusters across samples was correlated (Fig. 1F), finding that one particular cluster (cluster 6, n=116) behaved in a manner that was opposite to that of all other clusters.

Probing into the underlying commonalities between the genes that drove the clustering, we discovered that mean expression value for eight of the clusters (clusters 6-13) covaried with rosette size in both inbreds and hybrids. The most distinct cluster 6 showed a clear positive correlation, while the other clusters were negatively correlated with rosette size (Fig. 2A). While the general trend of correlation between gene expression and rosette size remained the same in F_1_s and the inbred parental lines, the shape of the correlation differed.

**Figure 2.**
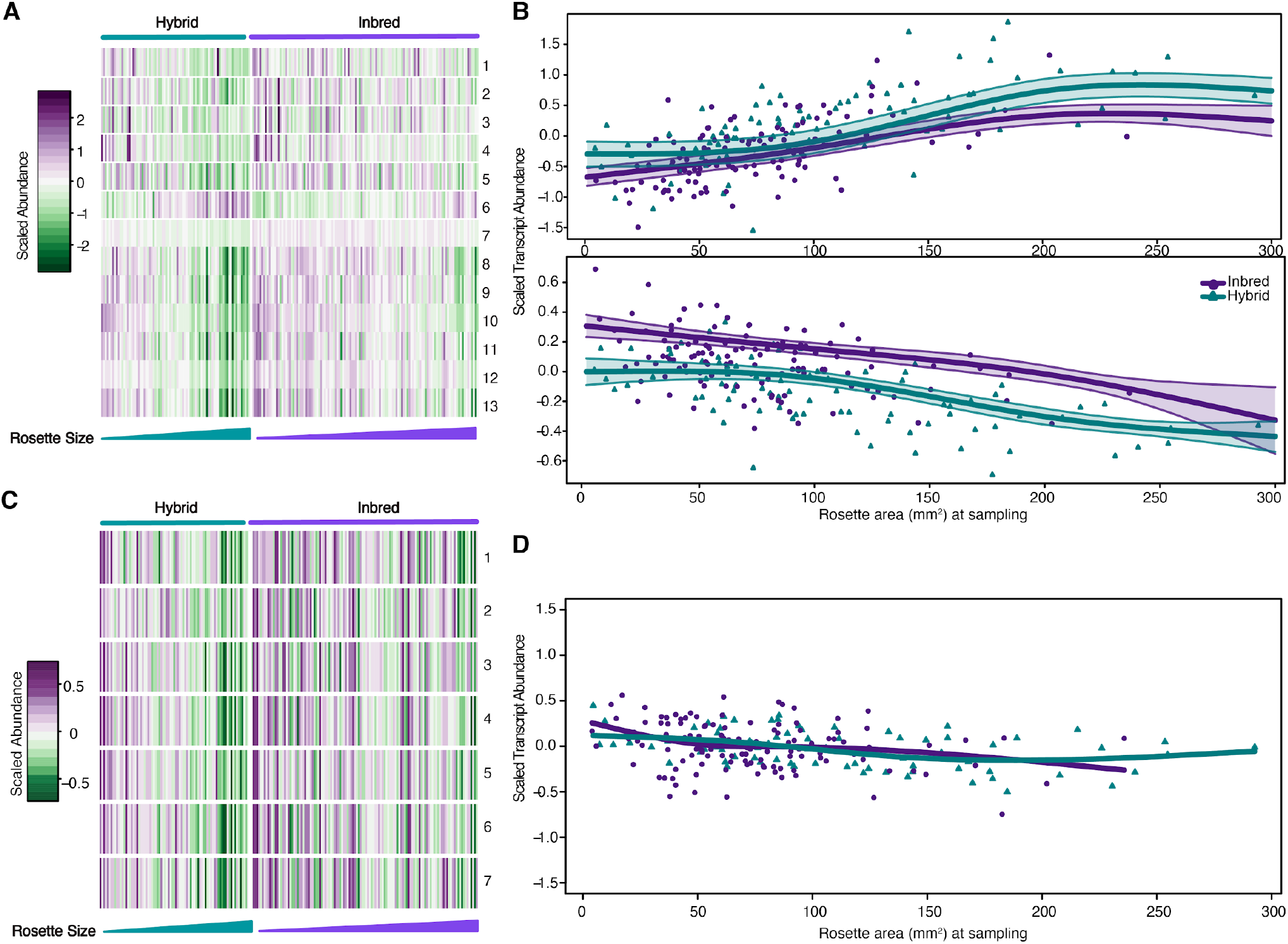
Non-additive gene expression level correlates with final rosette size. **A**. Heatmap showing the average expression of each non-additive gene cluster in each sample, sorted into F_1_s and inbred parental lines, and arranged by ascending final rosette size. **B**. Linear-Mixed-Model spline fitting of exemplary clusters. Top: cluster 6, bottom: cluster 7; points: cluster mean expression in each sample; shaded area: 95% Bayesian credible intervals. **C**. Heatmap of the average expression of gene clusters from 500 randomly sampled genes, arranged in the same order as in Fig. 2A. **D**. Randomly sampled genes, which show little expression-size covariation and no differences between inbreds and F_1_s.

To obtain further insight into the above observations, we formally investigated the relationship between average gene expression of each cluster and rosette size, by performing Bayesian linear-mixed-model spline fitting (Fig. 2B, Fig. S4, Methods). Clusters 1-5 showed no clear trend of association between expression and size, cluster 6 showed expression increasing in parallel with rosette size (Fig. 2B, top panel), and the remaining clusters showed expression decreasing with increasing rosette size. For these seven clusters, F_1_ hybrids tended to have lower expression than the inbreds across the entire range of rosette size (Fig. 2B, bottom panel). GO analysis did not indicate that clusters were specific for particular biological processes.

Therefore, a large number (n=1,420) of non-additively expressed genes showed covariation of expression level with rosette size. F_1_ hybrids systematically exhibited a shift towards either lower or higher expression levels in the direction consistent with the change in rosette size relative to inbred parental lines. The trend is unique for non-additive genes, as repetitive random subsets of the transcriptome showed neither profound covariation with size nor a systematic shift in the expression level in F_1_s (Fig. 2C-D).

### F_1_ exhibits robust growth advantage

That gene expression exhibited a systematic shift in the F_1_ hybrids, and that non-additive gene expression in the F_1_s covaried with rosette size prompted us to ask (i) whether degree of non-additive expression within individual parent-hybrid trios correlated with rosette size differences between the F_1_s and their parents, and (ii) whether a global perturbation to the plant’s developmental program would affect the F_1_s and the inbreds differently. To this end we conducted a second experiment in which we applied BTH (acibenzolar-S-methyl), an analog of the defense hormone salicylic acid (SA) (Fig. 3A, Methods), to 40 parent-F_1_ trios. Induction of pathogen defense was chosen as treatment because it causes morphological changes and at least sometimes extensive transcriptional reprogramming, as has been observed in some F_1_ hybrids of *A. thaliana (Bomblies et al. 2007)*.

**Figure 3.**
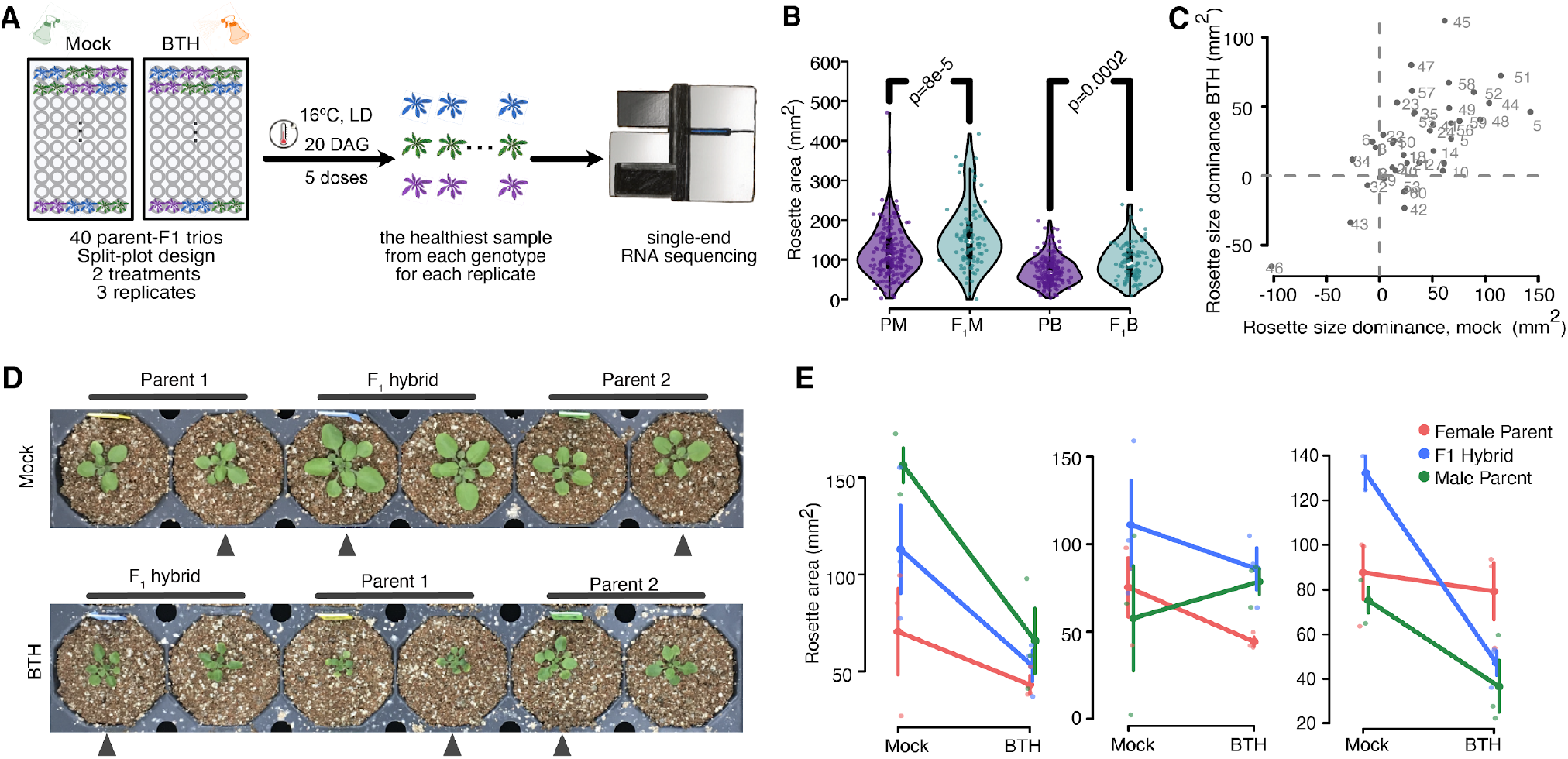
BTH treatment reduced rosette size in both inbreds and F_1_s. **A**. Experimental design. **B**. F_1_s maintained robust growth advantage despite the reduction in rosette size upon BTH treatment. PM: parent mock, F_1_M: F_1_ hybrid mock, PB: parent BTH treated, F_1_B: F_1_ BTH treated. **C**. Positive correlation between rosette size dominance under mock and BTH conditions. Numbered labels indicate the ID of the SHB2 trios. **D**. Typical rosette phenotype of a trio. **E**. Diverse response of three example trios to BTH treatment. Reaction norm lines connect the mean ± SD rosette area of each genotype under both treatments.

F_1_ hybrids are on average considerably larger than the inbred parents (inbred mock: 118.2±61.1 mm^2^, F_1_ mock: 157.1±79.6 mm^2^, inbred BTH: 71.6±35.5 mm^2^, F_1_ BTH: 94,3±44.3 mm^2^, Fig. 3B), consistent with our earlier experiment (Fig. S5). Neither the distribution of the rosettes of the parents nor those of the F_1_ plants was normal, with the F_1_ population having a significantly higher number of larger individuals (p=8e-5, two-tailed Kolmogorov-Smirnov).

In a comparison of randomly chosen trios, the F_1_ was almost twice as likely as one of the inbreds to be the larger individual (Cliff’s delta=0.31). BTH treatment significantly reduced plant size in both inbreds and hybrids (mock: 79.2±3.0 mm^2^, BTH: 39.5±4.2 mm^2^, One-way ANOVA, p<2×10^−16^, Fig. 3B, D, Fig. S6), and induced considerable variation in growth responses (Fig. 3E, Fig. S7). F_1_s exhibited stronger size reduction but remained to be more likely to be larger than either parent (p=0.0002, two-tailed Kolmogorov-Smirnov, Cliff’s delta=0.31). Most trios showed similar patterns in rosette size growth emergence after both mock- and BTH-treatment, with the majority of F_1_s remaining larger than the MPV (Fig. 3C).

Growth advantage is therefore prevalent in the F_1_ hybrids included in both of our experiments (Fig. 3B, Fig. S5), and is to a great extent robust to a perturbation of the developmental program. Although we cannot rule out that even stronger BTH treatment would eventually render hybrids smaller than the inbreds, it is, however, unlikely to occur in a natural setting, as our treatment already resulted in extremely dwarfed plants.

### Degree of non-additivity correlates with F_1_ growth advantage

Having established that non-additively expressed genes are systematically associated with plant size, and that F_1_ rosettes frequently exhibit positive size emergence, we investigated whether the degree of expression non-additivity in F_1_s may be associated with this phenotypic non-additivity. We focused on genes showing general response to BTH treatment (n=6,371) and asked whether deviations of F_1_ expression values from the MPV of any gene in individual trios exhibited correlation to the non-additivity in F_1_ rosette size. Clear correlations could be observed for many genes, which could be broadly categorized into (monotonic) positive, (monotonic) negative, or quadratic (Fig. 4A), while in some cases no correlation was observed. We defined 61 groups of genes that fell into these different categories by k-means clustering. To establish the significance of the size-expression correlation, we performed a Wilcoxon signed-rank test on the genes in each of the 61 clusters (Fig. 4B, Methods, Data S3). Some clusters shared similar relationships between the non-additivity in gene expression and rosette size, therefore we sorted the clusters further into 12 classes reflecting the pattern of correlation under mock and BTH treatment (e.g., “negative-negative” means negative correlation under both treatments, Fig. 4C, Methods). Correlation often changed in response to treatment, with the majority of genes exhibiting negative correlation with rosette size non-additivity under at least one condition, a trend that increased after BTH treatment (Fig. S8). This observation is consistent with your finding that negative dominance is more pervasively associated with rosette growth in *A. thaliana*.

**Figure 4.**
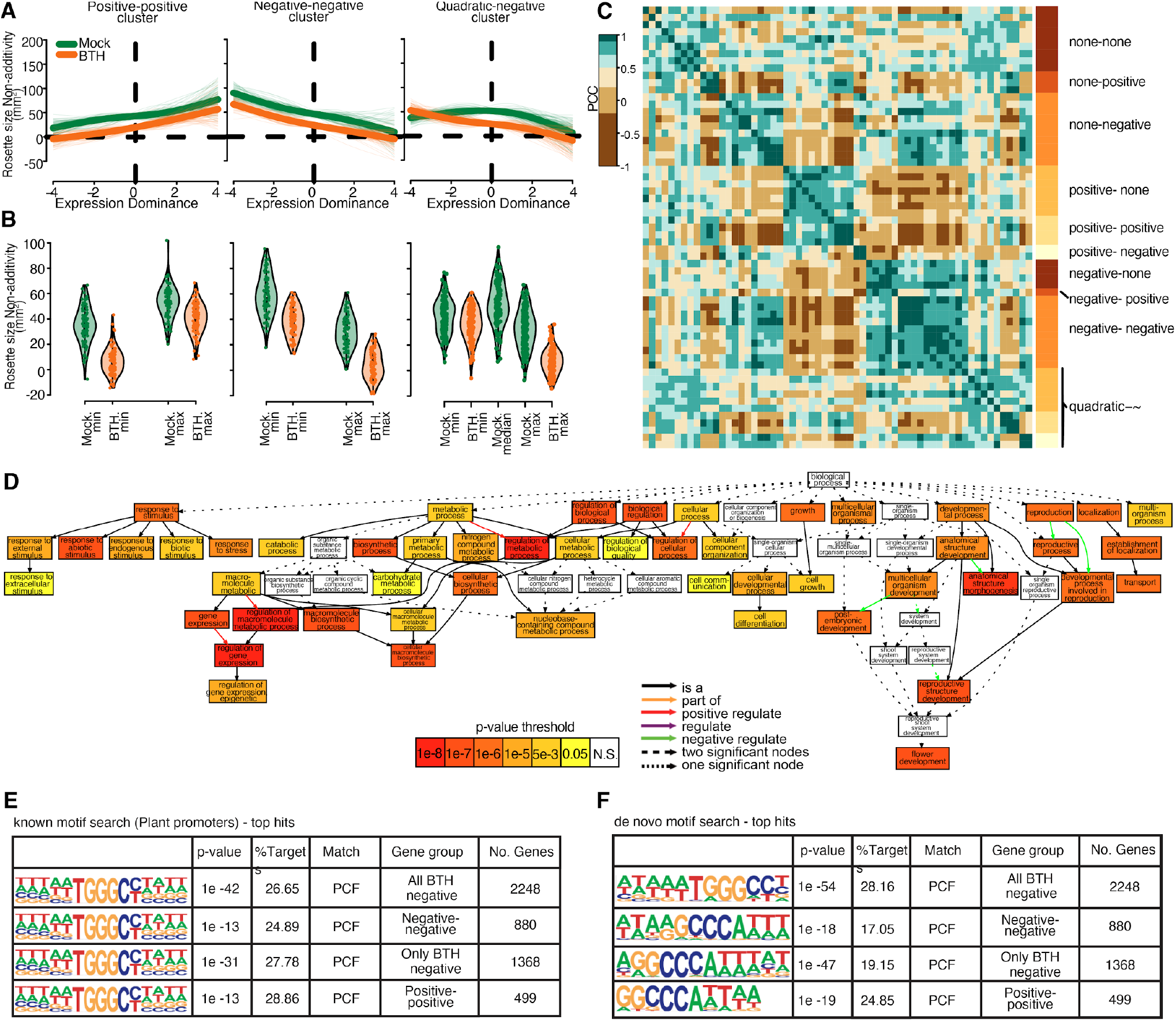
Genes whose degree of non-additive expression in trios correlates with hybrid performance. **A**. Exemplary clusters of “positive-positive” (left), “negative-negative” (middle), and “quadratic-negative” (right) genes. Thick solid line: spline fitting of the cluster means; thin lines: spline fitting of individual cluster members. **B**. Average biomass MPH for rosette samples with low- vs. high-expression of genes in the same clusters as in A. Each violin depicts the distribution of cluster gene expression averaged across the top and the bottom (and the middle for the quadratic relationship) deciles of samples. **C**. Pearson correlation coefficients (PCC) of all 61 clusters based on LMM-spline modeling. The clusters are further sorted into 12 classes labeled on the right according to the relationship between gene expression and biomass under mock or BTH treatment. **D**. GO enrichment for genes from the negative cluster; see Fig. S10 for more details. **E**. Regulatory regions of genes from both positive and negative clusters are enriched for a PCF binding motif. **F**. De novo motif search confirmed enrichment of the PCF motif.

The genes with negative size-expression correlation after BTH treatment (Data S4, “negative genes” hereafter) are enriched for GO terms “regulation of gene expression”, “floral organ development”, and “response to (abiotic) stimuli” (Fig. 4D, Fig. S9). Genes with positive correlation under both treatments (Data S5 “positive genes” hereafter) were moderately enriched for photosynthesis (Fig. S10). While it seems unlikely, we cannot exclude that the enrichment for abiotic response is due to our analytic focus on BTH-responsive genes, although a parallel GO enrichment test for all BTH-responsive genes did not return significant hits in any biological process. Together these results suggest that the F_1_ transcriptome is systematically repressed in reproductive growth and (abiotic) stress response functions while activated in photosynthesis. Such non-additive transcriptome signature is associated with the rosette growth advantage of the F_1_s, which is pervasive in our system.

To begin to discover potential regulatory mechanisms, we performed motif enrichment analysis among the heterosis associated genes (Methods). PROLIFERATING CELL FACTOR (PCF) and c-Myc transcription factor binding motifs are highly enriched in the promoters of negative genes (569/2,248 genes, p=10^−42^, and 433/2,248 genes, p=10^−21^, Fig. 4E, Fig. S11). PCF-binding motifs are also highly enriched in the positive genes (144/499 genes, p=10^−13^, Fig. 4E). These findings were corroborated by *de novo* motif searches (Fig. 4F). PCF/TCP proteins constitute a conserved plant-specific transcription factor family that includes several regulators of cell cycle, growth, and disease resistance (Gonzalez 2015; Li 2015). That both positive and negative genes were enriched for PCF motifs points to these factors as a potential central toggle for global re-modeling of hybrid transcriptomes.

Parenthetically, close to 900 additively expressed genes were called from the second experiment, and 300 of which (hereafter, common additive genes) overlapped with the first experiment (Fig. S12, Data S6, Methods). GO-term analysis of the common additive genes revealed enrichment in cell death- and stress response-related processes (Fig. S12), again confirming that these pathways are systematically buffered in F_1_s. As expected, the additive genes from the second experiment were more enriched for defense response than the genes from the first experiment without BTH treatment (Fig. S13).

### Dominant, not additive complementation underlies emergence of F_1_ growth advantage

We next asked whether the association between expression level and F_1_ growth has a common genetic basis. Hybrids offer a genomic playground where deleterious alleles from each of the parental genomes may be complemented, one of the leading hypotheses for growth advantage in hybrids. Deleterious alleles are expected to be segregating at lower frequency in the population, and those residing in the gene regulatory regions have been associated with mis-expression of genes in *cis* (Kremling et al. 2018). To gauge the effect of genetic complementation in the F_1_s, we turned to regulatory rare alleles, looking into the relationship between gene expression level and the average number of regulatory rare SNPs (within 1 kb upstream of genes), which are more likely to have a deleterious effect on gene expression than common SNPs, in both inbred parents and the F_1_s (Methods).

Regardless of the treatment, we observed on average a significantly higher rare allele counts upstream of the common additive genes associated with low expression ranks in inbred parents (Fig. 5A, Wilcoxon signed rank-sum test with Benjamin-Hochberg FDR: mock, α=1.1×10^−5^, BTH: α=1.1×10^−5^). The trend was moderate in the positive genes (Fig. 5B, mock: α=0.03, BTH: α=0.10) and in the opposite direction in the negative genes (Fig. 5C, mock: α=7×10^−6^, BTH: α=7×10^−7^). Trends in F_1_s were in all three gene groups consistent with what was seen in the parents (Fig. 5D-F, α=5.1×10^−5^, additive-mock; α=3.8×10^−4^, additive-BTH; α=0.005, positive-mock; α=0.23, positive BTH; α=3×10^−12^, negative-mock; α=3×10^−7^, negative-BTH). Note that for the non-additive genes, increased upstream rare-allele count was always associated with an expression pattern that is consistent with a smaller plant. Upstream rare-allele burden therefore tends to lead to more extreme, and deleterious expression of these genes, in parents and the hybrids alike.

**Figure 5.**
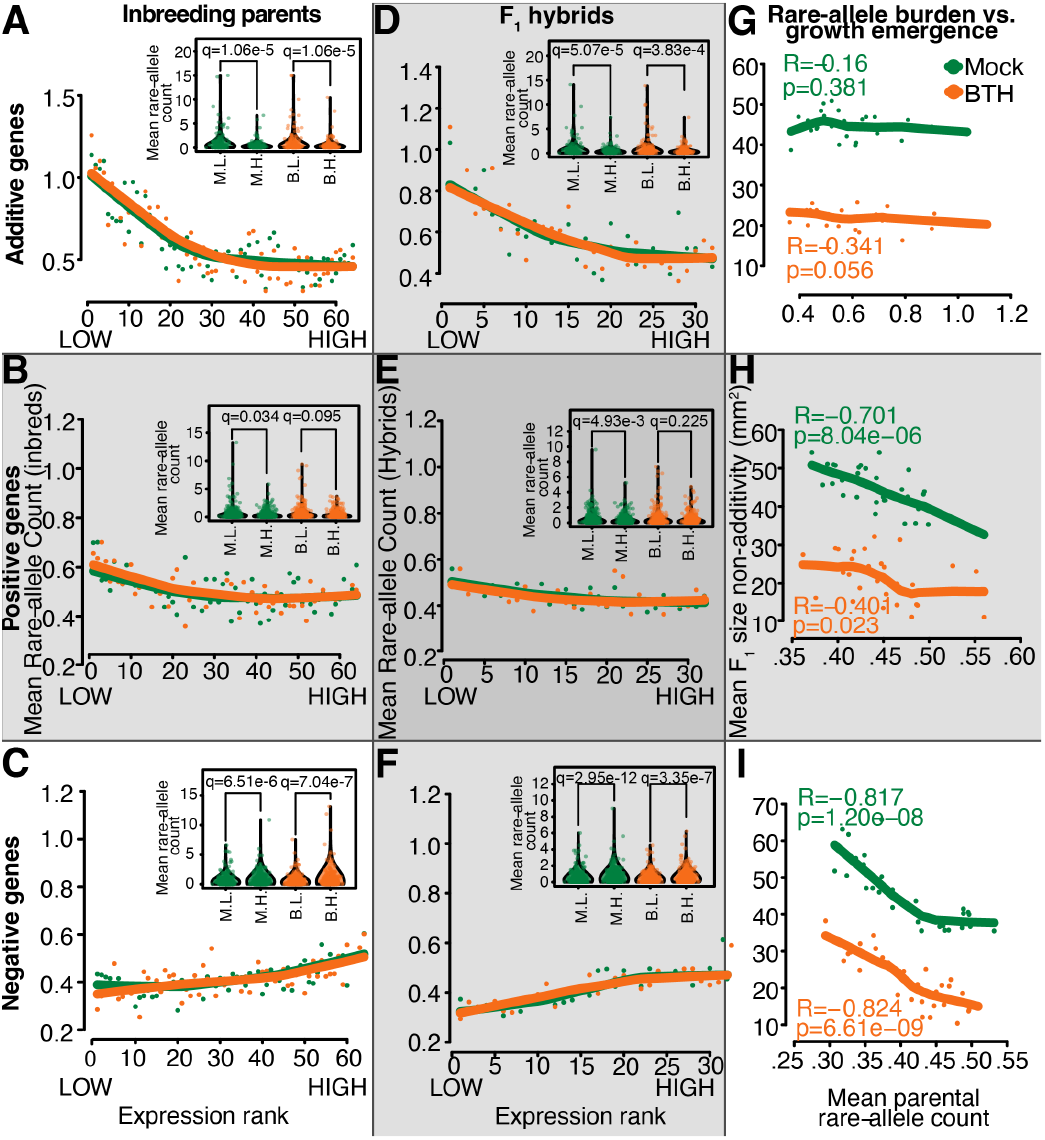
Rare allele burden affects gene expression and emergence of F_1_ growth advantage. **A-C**. Association between gene expression rank and upstream rare allele count of additive genes (A), positive genes (B), and negative genes in inbred parents (C). Samples were ranked by expression value for every gene within the gene list. Points: average upstream rare allele counts of all samples sharing the same rank; lines: LOWESS trend lines. Insets show the upstream rare-allele count of samples in the top (Mock 10, BTH10) and bottom decile (Mock 1, BTH 1) of expression ranks. **D-F**. Association between gene expression rank and upstream rare allele count of three gene lists in F_1_ hybrids. **G-I**. Association between rosette growth emergence and mean parental rare allele burden in additive (G), positive (H), and negative (I) genes. For each gene, F_1_s were ranked by average number of rare alleles in their parents. Points: average non-additivity in rosette size of all F_1_s sharing the same rank; lines: LOWESS trend lines

We next asked what the likely phenotypic consequence of the upstream rare-allele burden might be. Because non-additive gene expression varied less with rare-allele burden than additive gene expression (Fig. 5A, D), we expected these additive genes to be critical for F_1_ growth advantage. To our surprise, the number of rare alleles upstream of these common additive genes did not seem to be particularly relevant for rosette growth advantage in the F_1_s (Fig. 5G, Pearson correlation R=-0.16, p=0.38). Additive complementation may therefore be an inherent property of hybrids without directly influencing growth advantage. In contrast, the number of mean upstream rare alleles in both positive (Pearson correlation R=-0.701, p=8.04e-6) and negative (Pearson correlation R=-0.817, p=1.20e-8) genes is a strong negative predictor of rosette growth advantage in F_1_ hybrids (Fig. 5H-I, suggesting that growth advantage tends to be greater in hybrids derived from parents with fewer rare alleles in the upstream regions of these non-additively expressed genes. We conclude that dominant complementation contributes heavily to the F_1_ growth advantage observed in our system.

## Discussion

Most, if not all, of the growth and survival phenotypes of an organism are complex traits. Although the infinitesimal model predicts that non-additive effects contribute only minimally to most structural traits (Crow 2010), their impact is expected to be far greater in fitness-related traits (Merilä and Sheldon 1999). To date most of the methods for mapping and genomic selection are based on additive effects only (Varona et al. 2018). However, incorporating non-additive, *i*.*e*.,, dominant and epistatic, effects into quantitative genetics modeling can improve heritability estimates and accuracy of genomic prediction (Su et al. 2012; Wellmann and Bennewitz 2012; Kumar et al. 2015; Xiang et al. 2016). Limited empirical data exists for non-additive variance estimates, which varies greatly between organisms and can range from around 3-15% of total phenotypic variance in humans and animals (Misztal 1997; Zhu et al. 2015) to a third or more in plants (Kumar et al. 2015; Berguson et al. 2017; Calleja-Rodriguez et al. 2021). Adequate knowledge of non-additive genetic action is therefore of pivotal importance for a thorough understanding of the genetic architecture of complex traits, especially those that are fitness-related, and their evolutionary trajectory.

Motivated to better understand the prevalence and consequence of non-additive genetic action, we systematically compared the additivity and non-additivity in gene expression in *A. thaliana* F_1_ hybrids. Consistent with findings in other systems (Fujimoto et al. 2012; Bell et al. 2013), we found non-additivity to be prevalent. Moreover, there was a prominent association between non-additivity and biomass on a cross-population scale, lending support to the theory of directional dominance being an underlying factor of emergent phenotypes in hybrids (Falconer and Mackay 1996).

Hybrid advantage was common in our F_1_ collection, consistent with inferences from smaller sets of *A. thaliana* F_1_ hybrids (Seymour et al. 2016; Oakley et al. 2019). In our system, hybrid advantage seems to be driven in large part by non-additive gene expression. For many genes, when parent-hybrid trios are considered, the degree of non-additivity in F1 expression is either positively or (more frequently) negatively associated with hybrid performance. GO enrichment points to repression of reproductive development and abiotic stress response, as well as increased photosynthesis as factors that support more robust growth in F1 hybrids. The rewiring of transcriptional networks is likely to occur in *trans*, as binding motifs for a small number of transcription factors were enriched in the promoters of the focal genes.

In contrast, additive (i.e., near-MPV) expression itself does not correlate with the larger size of F1 hybrids, hence probably does not directly contribute to hybrid advantage. The observation is consistent with the notion that additive variance is quickly driven to fixation in fitness traits (Merilä and Sheldon 1999). Additive expression appears to be an intrinsic property for certain genes, mainly enriched in cell-death and stress-response pathways, in F1 hybrids, largely independent of specific parental combinations. Whether tighter control of biotic defense responses capacitates hybrid advantage requires further investigation.

It has long been postulated that heterosis is driven by directional dominance (Falconer and Mackay 1996). It is therefore noteworthy that we observed a consistent species-wide trend of non-additive expression leading to the relative gene-expression level in F1 hybrids being more similar to the low-expression parents. With the proviso that we cannot rule out that the absolute transcript abundance of these genes were in fact increasing, our measurements do indicate a relative decrease in cellular transcript concentration, which could have consequences for molecular interactions (Gao et al. 2021; Koşar and Erbaş 2022). The abundance of up-stream rare alleles in these non-additively expressed genes further suggests that the change of expression had a genetic basis, as opposed to being merely a consequence of a common phenotype.

Another study recently reported high-parent expression of hub genes from regulatory networks of photosynthesis and cell cycles during early shoot development to be associated with a high degree of growth advantage in one specific F_1_ hybrid (Liu et al. 2021). We similarly found above-MPV expression of photosynthetic function in the later phase of vegetative growth to be positively correlated with increased growth. Both studies also agree in low-parent gene expression being common in F_1_ hybrids during the later stage of vegetative growth.

We observed putative deleterious effects (*i*.*e*., strong deviation of gene expression from that of the population mean) of rare alleles in upstream regulatory sequences, primarily in inbred parental accessions, and to a lesser degree in F1 hybrids, suggesting that rare-allele burden has common effects in inbreds and hybrids alike. The correlation between rare-allele burden and gene expression is more pronounced in additively than non-additively expressed genes. However, while (the milder) rare-allele burden in non-additive genes is strongly associated with phenotypic consequences in F1 hybrids, this is not reflected in the behavior of the additive genes. We conclude that concerted non-additive gene expression, rather than canalization via additive gene expression, is a main driver of growth advantage in *A. thaliana* F_1_ hybrids.

The use of RNA-seq enabled the parallel comparison of thousands of gene expression traits, all quantified against the same scale. With this, we could define both “additive” and “non-additive” genes by their expression at the population level, largely circumventing idiosyncratic behavior of genes in a specific trio, enhancing our confidence that our observations can be generalized across the species. By associating transcriptome changes and plant growth, we were also able to characterize growth as a high-dimensional phenotype, with under-dominance as the predominant type of gene action associated with growth advantage in F_1_ hybrids. Our work provides another step towards understanding molecular mechanisms and evolutionary forces that lead to dominance complementation of rare regulatory alleles.

## Materials and Methods

### Generation of genetic resource

Accessions covering the entire species range were chosen from the *A. thaliana* 1001 Genomes Project (1001 Genomes Consortium). F_1_ hybrids used in this study were generated via random crosses, either by randomly crossing individual accessions that reached flowering stage at the same time (SHB1) (Vasseur et al. 2019), or according to a pre-generated randomized crossing scheme after subjecting seedlings to a saturating (12 weeks) vernalization under 4°C short-day conditions (SD, 8/16 hr photoperiod) to synchronize flowering (SHB2).

### Experimental design

The first experiment (Fig. 1A, “SHB1”) initially included 101 parent-F_1_ trios of altogether 286 distinct genotypes. Single plants were grown following an incomplete randomized block design, with each tray as a block within which each genotype was sown as an adjacent pair. The second experiment (Fig. 3A, “SHB2”) included 40 parent-F_1_ trios. Single plants were grown in triplicates, following a split-block design where each block held 10 parent-F_1_ trios with each row consisting of one trio with duplicates of plants in adjacent pots. The trios within each block, and the relative positions of genotypes within each trio were randomized. Plants were subjected to either a mock or an artificial defense hormone treatment (see below). After accounting for germination, survival, and initial filtering of RNA-seq outputs, 82 hybrids and 124 inbred parents in SHB1, and 32 trios in SHB2 remained for downstream analyses.

To minimize circadian bias, sowing for both experiments was scheduled in batches to ensure that harvesting could be finished within a 30-minute window at the same hour for several consecutive days. At 21 days after sowing, the healthiest appearing plant of each genotype was used for RNA-seq, to ensure any sampling bias is systematically towards the same direction for both inbreds and hybrids. Meanwhile rosette size measurements were obtained for the same individual plants for which RNA-seq were performed.

For a list of genotypes analyzed in both experiments, see Data S1.

### Plant culture, treatment, and sampling

Single plants were grown in a 1:1 mixture of calcined clay media (Diamond Pro, Arlington, TX, USA) and vermiculite (Floragard, Oldenburg, Germany) supplemented with liquid growth media (Conn et al. 2013). Plants were not vernalized, to ensure that they remained in the vegetative growth phase. As a proxy of vegetative biomass (ref. (Vasseur et al. 2018) and Fig. S1), the rosette area of 21-day old plants was measured. The full rosettes grown under 16°C long-day (LD,16/8 photoperiod) were harvested and flash-frozen at 21 day-after-germination (DAG).

Defense hormone treatment started 14 days after sowing. An analog of the defense hormone salicylic acid (SA), BTH (acibenzolar-S-methyl, Sigma-Aldrich) was used, with optimal dose and treatment scheme of the defense hormone that had been established in a pilot experiment on multiple accessions (Fig. S2). Each 10×6-pot-tray block was the unit of treatment and plants were treated by topical spraying every other day with either a mock solution (20 mL; ddH_2_O, 0.1% DMSO, 0.006% Silwet) or a BTH solution (20 mL; 100 mM acibenzolar-S-methyl, 0.1% DMSO, 0.006% Silwet), and covered for 1 hour with transparent plastic lids after spraying. A total of five treatments were administered. Full rosettes were harvested and flash frozen at 21 DAG.

### Growth analysis

Plant growth was monitored by daily image capture from the top of the trays using the RAPA system (Capovilla et al. 2018). Rosette areas were acquired by automatic image segmentation and counting of green pixels, supplemented with manual curation. The rosette size estimates were then converted from pixel counts to mm^2^ by multiplication with a calibration factor.

### RT-qPCR

To establish defense hormone treatment and dosage, the effect of salicylic acid and BTH application was tested by treating 18 accessions with Mock (ddH_2_O, 0.1% DMSO, 0.006% Silwet), 350 mM SA and 100 mM BTH in 3 replicates, each with duplicated plants for phenotyping and qPCR. After 5 treatments the rosettes were harvested in one set of plants to compare their sizes, while the other set of plants were used for qPCR to compare the effect of 350 mM SA and 100 mM BTH treatments. Specifically, RNA was extracted and reverse transcribed. qPCR was performed using SYBER green (Thermo Scientific Maxima SYBR Green qPCR Master Mix (2x)) and primers for *ACTIN2, UBC21, PR1* and *NPR1*. Normalization across plates was performed using the same set of samples featured on all plates. The data were analyzed by calculating ΔΔCq (Fig. S14).

### RNA-seq

RNA-seq libraries were constructed as described (Cambiagno et al. 2021), using 750 ng total RNA from full rosettes as input. All libraries, each carrying a unique barcode combination were pooled and sequenced in multiple single-end lanes on an Illumina HiSeq 3000 platform for a target coverage of 5M reads per sample.

### RNA-seq read mapping and post-processing

FASTQ files from multiple lanes were merged and mapped to TAIR10 transcriptome using *RSEM* (*bowtie2*) with default parameters. Libraries with more than 8M mapped reads were subsampled to 8M with *seqtk* prior to mapping. Transcripts mapped to chloroplast, mitochondria, rDNA clusters, transposable elements (TEs) and pseudogenes, as well as transcripts with effective length less than 150 nt were removed from the raw *RSEM* count file. TPM (transcripts per million) counts were then re-estimated for the rest of the genes. Libraries with fewer than 2M mapped reads and those identified as extreme outliers following a principal-component analysis (PCA) of whole-transcriptome log_2_ (TPM) values were excluded from further analysis. Gene lists were further filtered for average transcript abundance (trimmed mean of log_2_ (TPM)>0.3) and coefficient of variance >0.15.

### Additive gene calling

For SHB1 data, MPV of each gene was calculated for all complete Parent-F_1_ trios by taking the arithmetic mean of the parental log_2_ (TPM). Linear regression was then performed between corresponding F_1_ expression value and the MPV. For SHB2, a linear-mixed model was used to correct for treatment and batch effects. Genes were filtered for regression coefficient >0.5 and R^2^ >0.4 for SHB1, and regression coefficient >0.4 and sigma <0.6 for SHB2. All thresholds were determined by quantiles. Genes called in both SHB1 and SHB2 were taken as common additive genes.

### Non-additive gene calling

With SHB1, a population-wide MPV distribution was established for each gene by calculating arithmetic means of log_2_ (TPM) between all possible pairwise combinations of inbred accessions. Two-sided Kolmogorov-Smirnov test was performed per gene to test if the log_2_ (TPM) from the F_1_ hybrids were drawn from the MPV distribution. Genes with q<0.001 (Benjamin-Hochberg FDR) were considered as non-additively expressed genes.

### Bayesian modelling of non-additive expression and plant size

non-additive genes from SHB1 were clustered by K-means, with the optimal K determined by the elbow method. A linear-mixed-model (LMM) spline was fitted using the *lme4* package(Bates et al. 2015) in R (ref. (R Core Team 2021)) for gene expression:

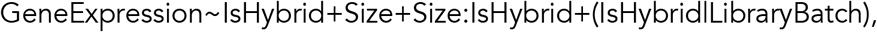

in which *GeneExpression* is the z-scaled log_2_TPM, *IsHybrid* is a binary code of the hybrid/inbred identity, *Size* is the z-score of rosette size at sampling. Natural cubic splines were modeled for *Size* and *Size-IsHybrid* interaction. The 95% credible intervals for the parameter estimates were established with 10,000 iterations of Bayesian simulation using the *arm* package(Andrew Gelman, Yu-Sung Su, Masanao Yajima, Jennifer Hill, Maria Grazia Pittau, Jouni Kerman, Tian Zheng, Vincent Dorie 2021).

### BTH responsive genes

The effect of BTH treatment on gene expression was identified by LMM:

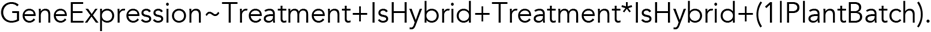

To establish a significant threshold, 10,000 permutations were performed for each gene, and the empirical p-value was corrected with Benjamin-Hochberg FDR. Genes with q<0.001 were kept as BTH responsive genes (n=8,797) and examined for their size-MPH correlation.

### Expression-plant size MPH correlation

Expression-MPH and size-MPH were calculated per trio by calculating the per-gene expression and rosette area difference between F_1_ and the MPV in corresponding treatments and replicates. Size MPH-to-expression MPH regression spline was acquired separately for both treatments. An initial round of K-means clustering was performed on the resulting spline coefficients, with the optimal K determined as the division with the highest Dunn index which allows no more than 25% of the clusters carrying less than 5% of the genes. Resulting clusters were inspected and removed if size and expression MPH do not co-vary. The remaining genes (n=6,371) were re-clustered with the same criteria, and the resulting clusters were manually sorted based on size-expression covariation into 12 general categories (Data S3, Fig. 4D, Fig. S9).

### Size-MPH covariation test

To establish the significance of the size-expression correlation, we performed a Wilcoxon signed-rank test on the genes in each of the 61 clusters (Fig. 4B). For each “none” cluster, each gene within the cluster was used as a data point, and the mean rosette size of 4 plants having the lowest and the highest expression MPH of the gene was calculated. The average rosette size corresponding to the two extremes of expression MPH were then compared with a two-sided Wilcoxon signed-rank test. Likewise, for “positive” and “negative” clusters, one-sided tests were used to test for significant differences between average rosette size corresponding to the two extremes of expression MPH within individual clusters. For “quadratic” clusters, separate one-sided tests were performed comparing the samples with extreme expression MPH against those with median expression MPH. Bonferroni correction was used to control for multiple hypothesis testing. The test revealed that our sorting procedure erred on the conservative side: while the top and bottom deciles were significantly different for 17 clusters assigned to the “none” category, only 6 of the “none” clusters were misassigned as “positive”, and none were misassigned as “negative” (Bonferroni corrected **α**<0.001, Data S3). The evidence for truly quadratic correlations was less clear.

### GO enrichment

GO enrichment was performed using the *Agrigo* v2 platform(Tian et al. 2017), with all gene IDs that passed our initial filtering (n=14,067, TAIR10 annotation) as background against plant GOslim database. Fisher’s Exact Test was used, and the enrichment p-value was corrected using Yekutieli FDR. The enrichment results were visualized with the built-in DAG-drawer of *Agrigo* v2.

### Rare allele analysis

Rare (MAF<0.05), biallelic SNPs 1 kb upstream of gene features were subset from the SNP annotations of the 1001 Genomes Project (1001 Genomes Consortium. Electronic address: and 1001 Genomes Consortium 2016). Genotype information at these SNPs were acquired for the accessions used in SHB2, and the sum of these rare SNPs upstream of each gene were calculated per accession. For F_1_ hybrids, upstream rare-allele count was determined by the mean of the rare-allele counts of both parents.

Samples, separated by inbred parents/F_1_ hybrids and with/without BTH treatment, were ranked for their expression values for each gene within a gene list of interest. For each given rank, a gene-list mean upstream rare-allele count was acquired by averaging across all samples received the same rank in any of the genes within the gene list. Relationships between gene-list mean upstream rare-allele count and expression rank were examined by LOWESS regression and tested with Wilcoxon signed rank sum test between the top and bottom decile of the expression rank.

Likewise, an average rosette-size MPH for each rank was calculated for the gene list, and the Pearson correlation was acquired between average rosette-size MPH and upstream rare-allele count.

### Motif enrichment and *de novo* motif finding

Motif enrichment and *de novo* motif finding was carried out using *HOMER* v4.10.4 (ref. (Heinz et al. 2010)) with TAIR10 reference genome and gene annotation. For every set of candidate genes, genomic sequences 1 kb upstream from the transcription start site (TSS) and 1 kb downstream from the transcription termination site (TTS) were indexed from the strand-specific gene coordinates. Both assays were performed by using the *findMotifsGenome*.*pl* function and *HOMER*’s in-built plant promoter motif database as reference:

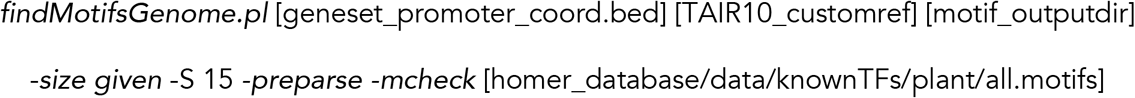

The top known motif hit (in all cases p≤10^−10^) from each candidate set was then used for a second motif enrichment step, where the promoters and downstream sequences were searched for the significant motif using *HOMER*’s *annotatePeaks*.*pl* function:

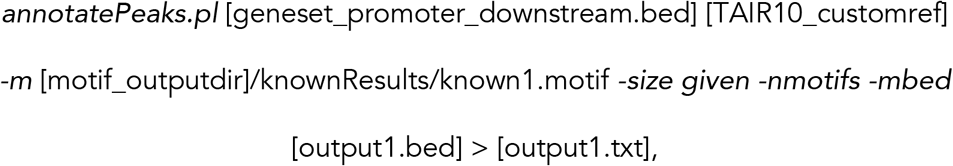

to generate corresponding genomic coordinates, and subsequently associated back to the genes containing the motif of interest in their regulatory regions (*bedtools* v2.26.0 intersect)(Quinlan and Hall 2010).

## Acknowledgements

We thank members of the Weigel lab for assistance with sample collection, and James Birchler, Chang Liu, François Vasseur, Ulrich Lutz, Rebecca Schwab, and Yiliang Ding for helpful discussions. This work was supported by the Max Planck Society.

## Author Contributions

Conceptualization – WY, DW; Methodology – WY, IB; Experiment – WY, FB, SS, RK; Data analysis – WY, FB, TS; Writing – original draft: WY, FB; Writing – review & editing – WY, DW.

### Data and Material Availability

Raw sequencing data is available at the ENA under the accession ERA9420648 and ERA9420737. Code to generate the results is available at: https://github.com/weigelworld/SigHeterosis.

## Supplementary Data Sets

**DataS1**.xlsx Sequenced and analyzed genotypes from SHB1 and SHB2

**DataS2**.txt Non-additive genes called in SHB1, and their cluster identity

**DataS3**.xlsx Wilcoxon rank sum test on the size-expression MPH spline clusters

**DataS4**.txt Genes negatively associated with heterosis under BTH or mock + BTH conditions (“negative genes”)

**DataS5**.txt Genes positively associated with heterosis under both mock and BTH conditions (“positive genes”)

**DataS6**.txt 300 common additive genes

## Supplementary Figures

**Figure S1.**
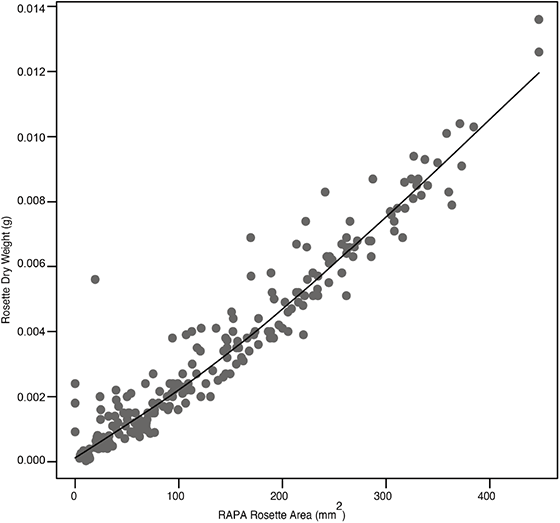
Rosette area serves as a good predictor of rosette biomass. Pearson correlation coefficient (R= 0.96, p<2.2e-16) of biomass (g) with rosette area (mm^2^) as measured on the RAPA system at 21 days after sowing of 221 individuals of mixed genotypes (grey dots), and LOWESS trendline.

**Figure S2.**
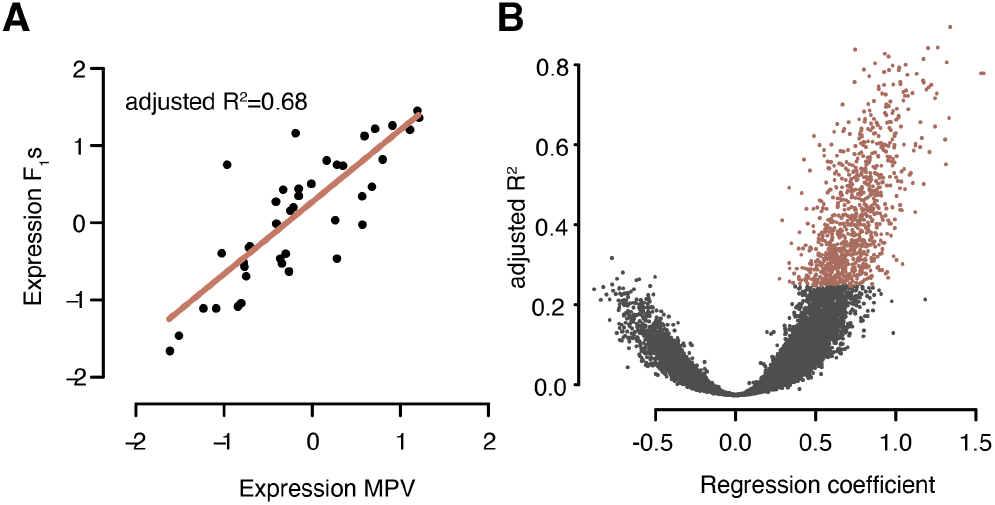
Linear-model-based additive gene calling. A. One representative additive gene (At5G43740, a CC-NBS-LRR class disease resistance protein, Non-additive rank: 16,574) whose expression in F_1_ hybrids closely correlates with calculated mid-parent value, both showed as z-score. The red line shows the regression line. B. Volcano plot of R^2^ value of each gene plotted against the corresponding linear-regression coefficient. Red dots show genes that passed the filtering threshold.

**Figure S3.**
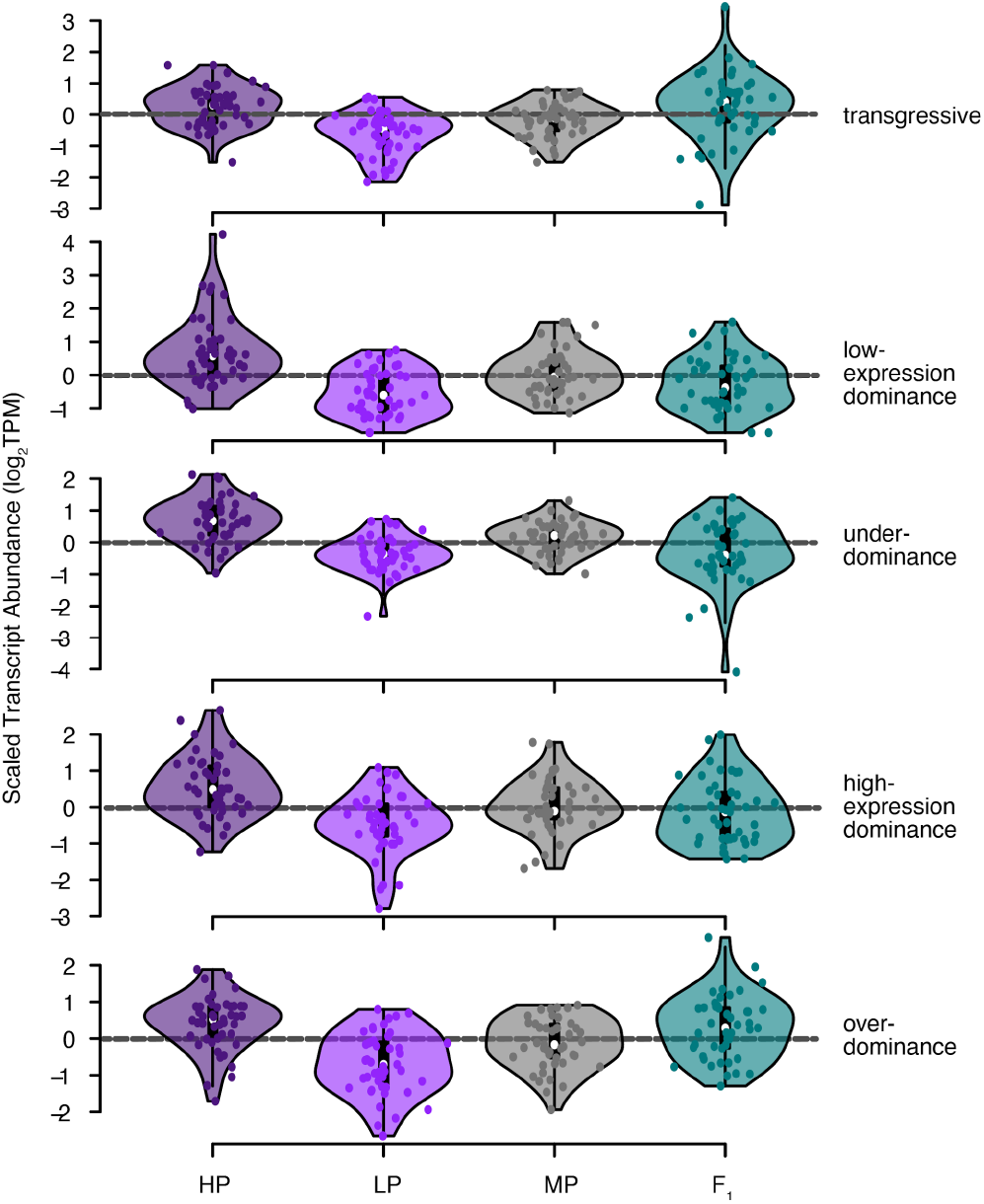
Examples showing that non-additive genes exhibit various forms of dominance in F_1_s. From top: AT5G07960, AT3G61170, AT2G17670, AT2G27510, AT4G12970.

**Figure S4.**
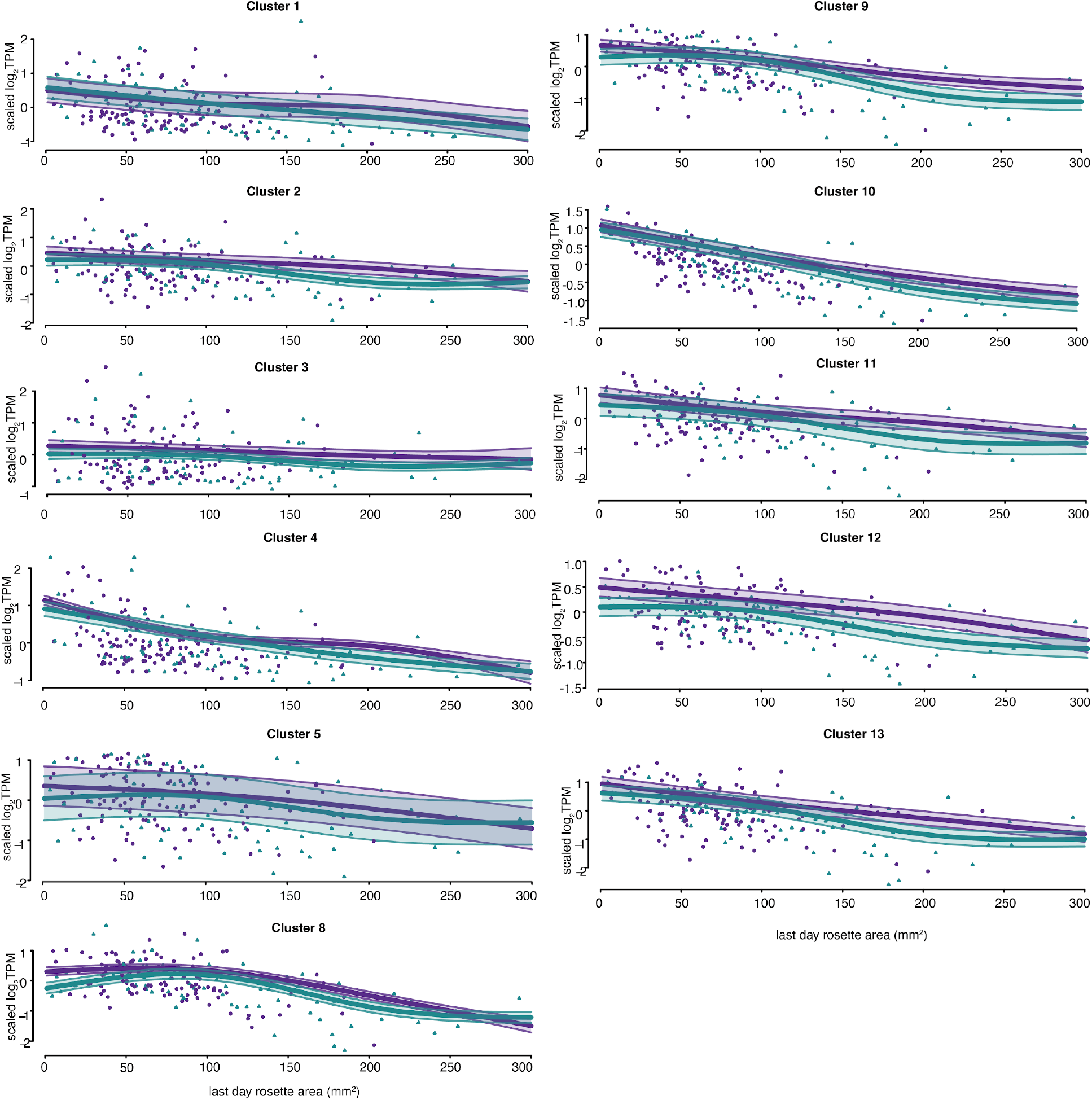
BTH treatment reduced rosette size in both inbreds and F_1_s. LMM spline fitting of cluster mean expression levels with 95% Bayesian credible intervals for non-additive gene clusters not shown in Fig. 3. Mean expression level across all genes within a cluster for inbred parents (purple dots) and F_1_ hybrids (turquoise dots) against last day rosette area (mm^2^) were plotted. Clusters 1-5 showed little rosette size-expression level association, while clusters 8-13 showed monotonic decrease of cluster mean expression level with increased plant size. While F_1_ hybrids exhibited the same trend as the inbred parents, the mean expression levels are consistently lower in F_1_ hybrids across the entire rosette size range for cluster 8-13.

**Figure S5.**
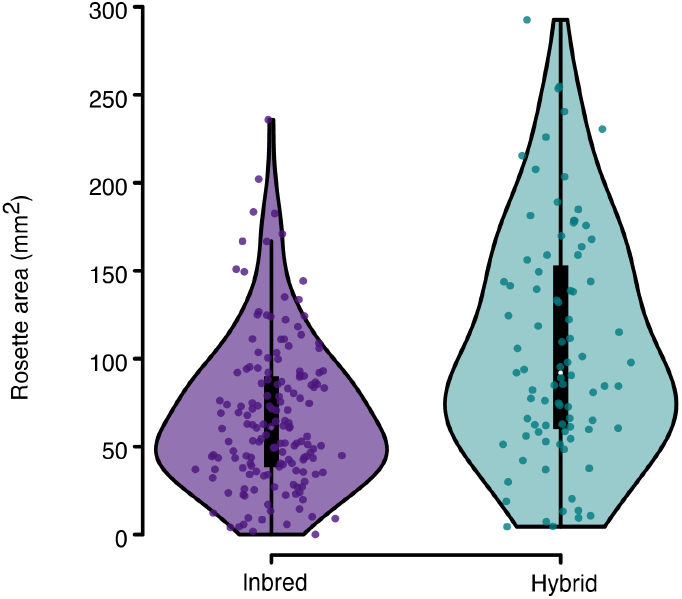
Rosette size in inbred parental lines and F_1_ hybrids. The F_1_ population has significantly more large individuals than the inbred population (p=2×10^−5^, Two-tailed Kolmogorov-Smirnov). F_1_s: 107.1±66.1 mm^2^, n=82; parents: 67.2±42.5 mm^2^, n=124. In a comparison of randomly chosen F_1_s and inbreds, the F_1_ hybrid was twice as likely than the inbred to be the larger individual (Cliff’s delta=0.33).

**Figure S6.**
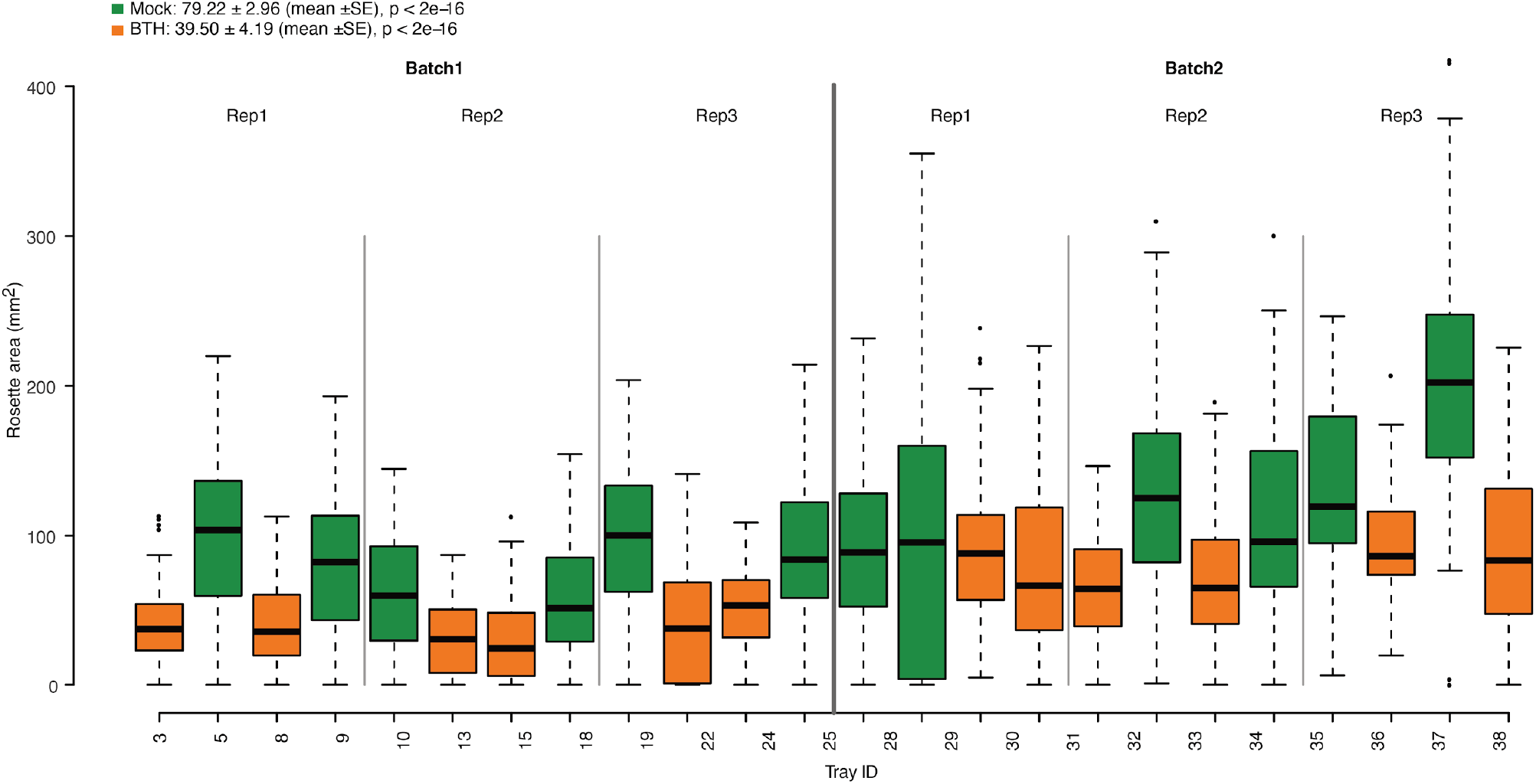
Consistent and significant reduction in rosette area by BTH in both batches of the SHB2 experiment. Each tray was a treatment unit. Shown are boxplots of rosette area (mm^2^) in each tray with mock (green) or BTH treatment (orange). Only trays with parent-hybrid trios further used for transcriptome analysis are included. The rosette area after treatment with BTH was reduced compared to mock treatment (2-way nested ANOVA, mock: 79.2 ± 3.0 mm^2^, BTH treatment: 39.5 ± 4.2 mm^2^; p < 10^−16^).

**Figure S7.**
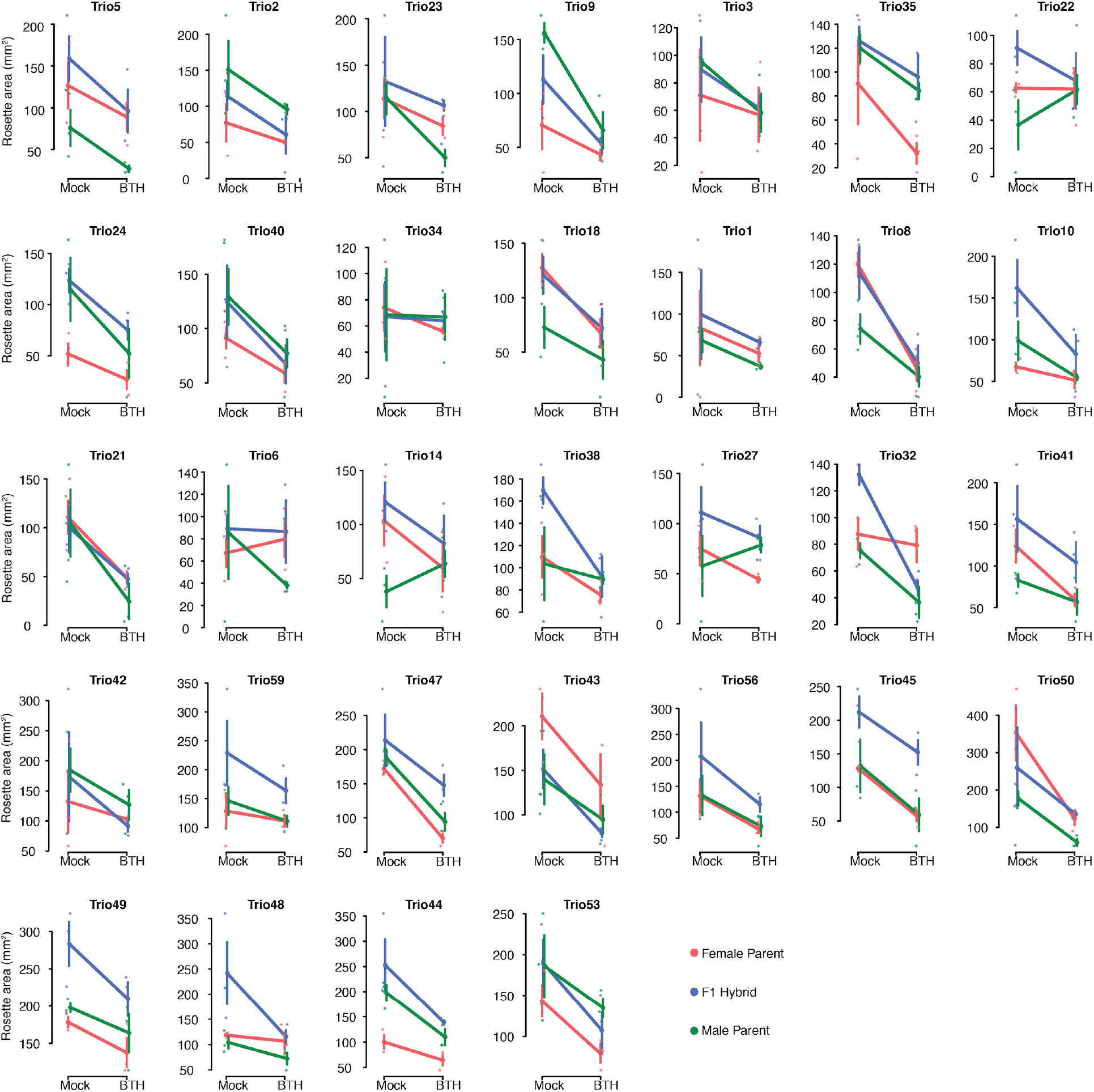
Reaction norm of rosette area (mm^2^) after mock and BTH treatments for all inbred parents and F_1_ hybrid trios. Solid lines connected the mean rosette area under the two treatments, with the dots illustrating individual rosette area of each plant and the error bars showing standard deviation of the biological replicates.

**Figure. S8.**
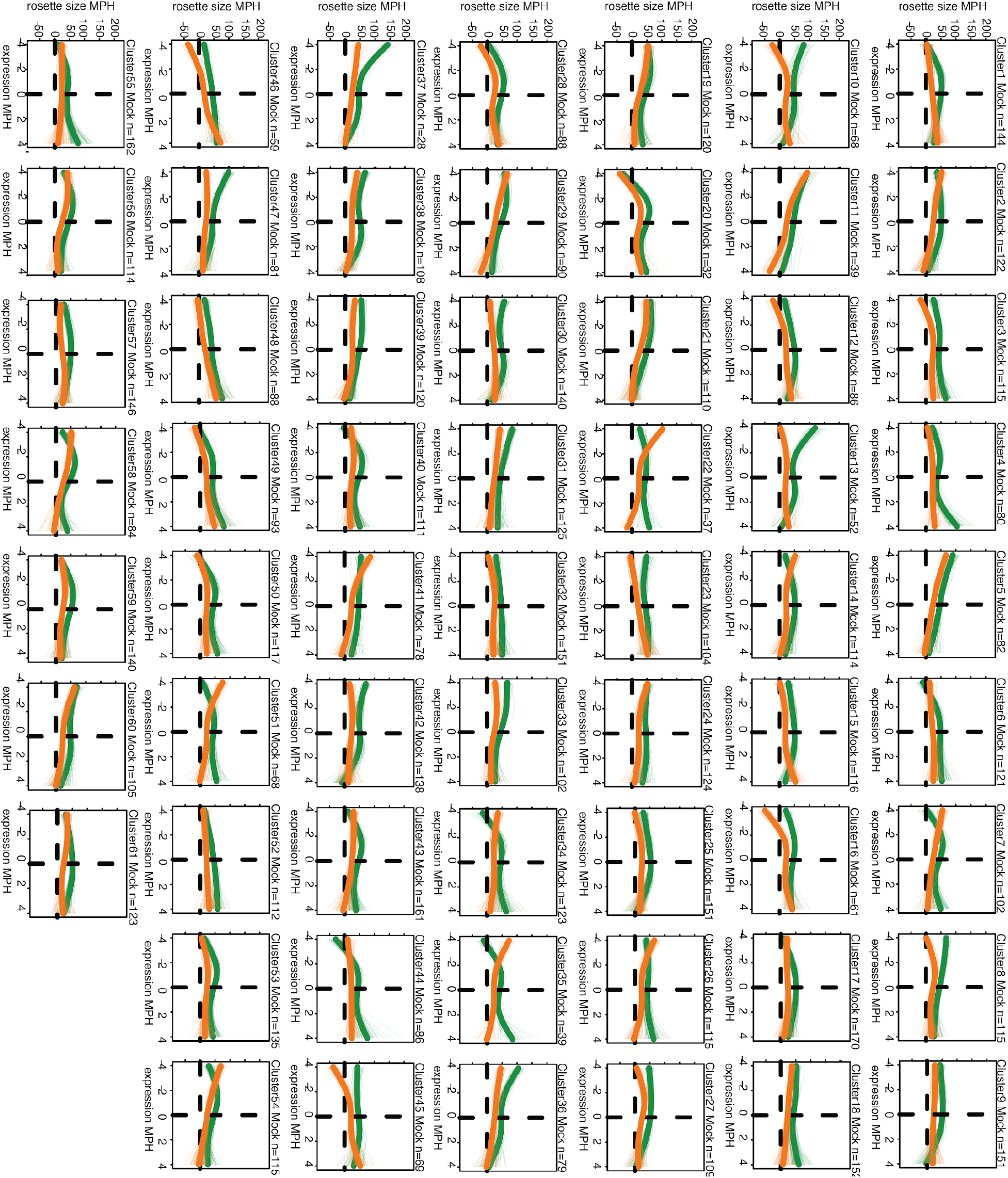
BTH-responsive genes sorted into 61 clusters. Genes were sorted based on spline regression of rosette area MPH (mm^2^) to their expression MPH across all trios. Shown are a graphic representation of the clusters. Each thin line represents a gene, and the thick line represents cluster mean (green: mock, orange: BTH). The 61 clusters were subsequently sorted into 12 general categories based on the regression trends in mock and BTH treatments.

**Figure S9.**
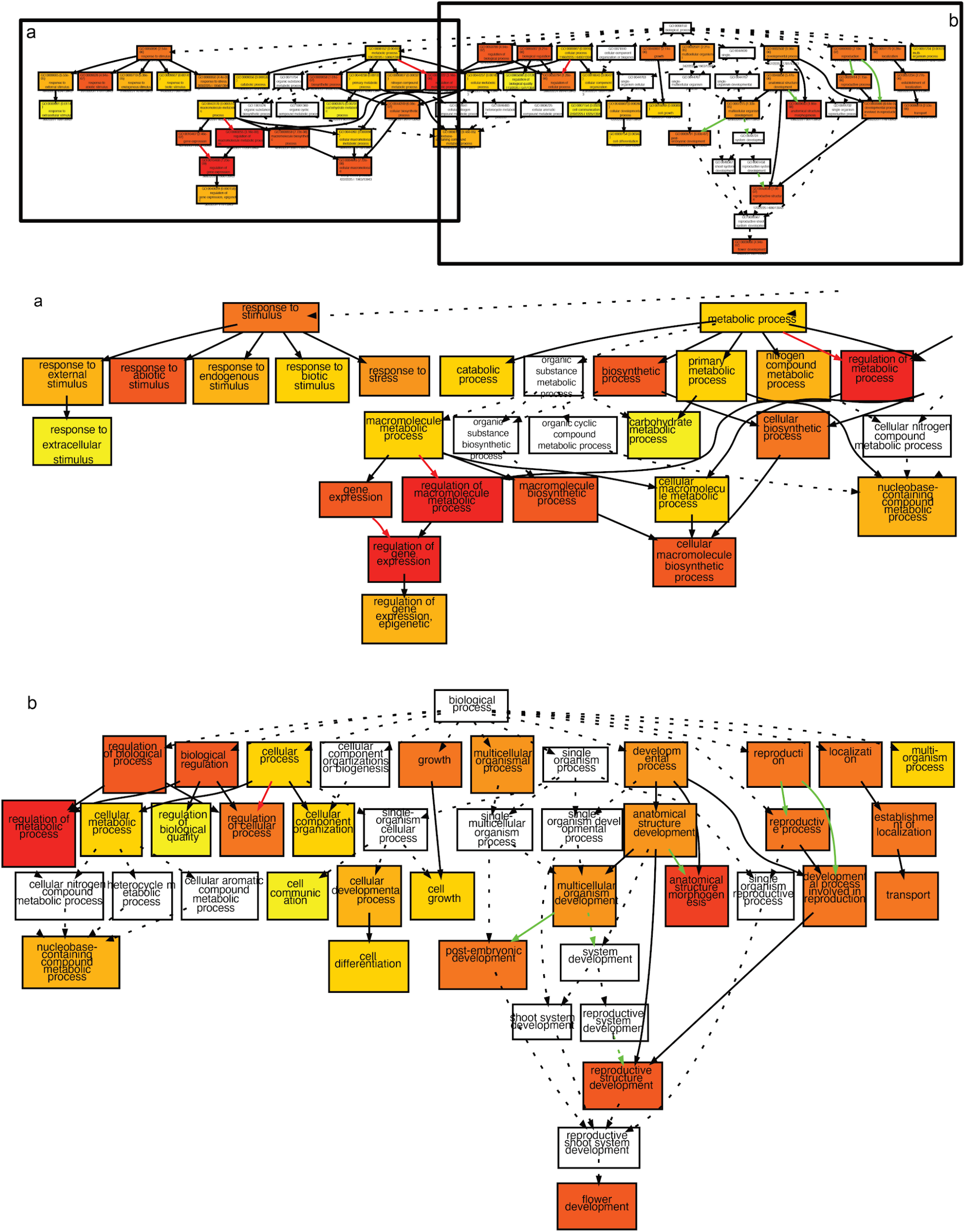
GO enrichment for the negative genes. Enlarged rendering of Fig. 4D. The diagram was split into two parts (a and b) for improved legibility.

**Figure S10.**
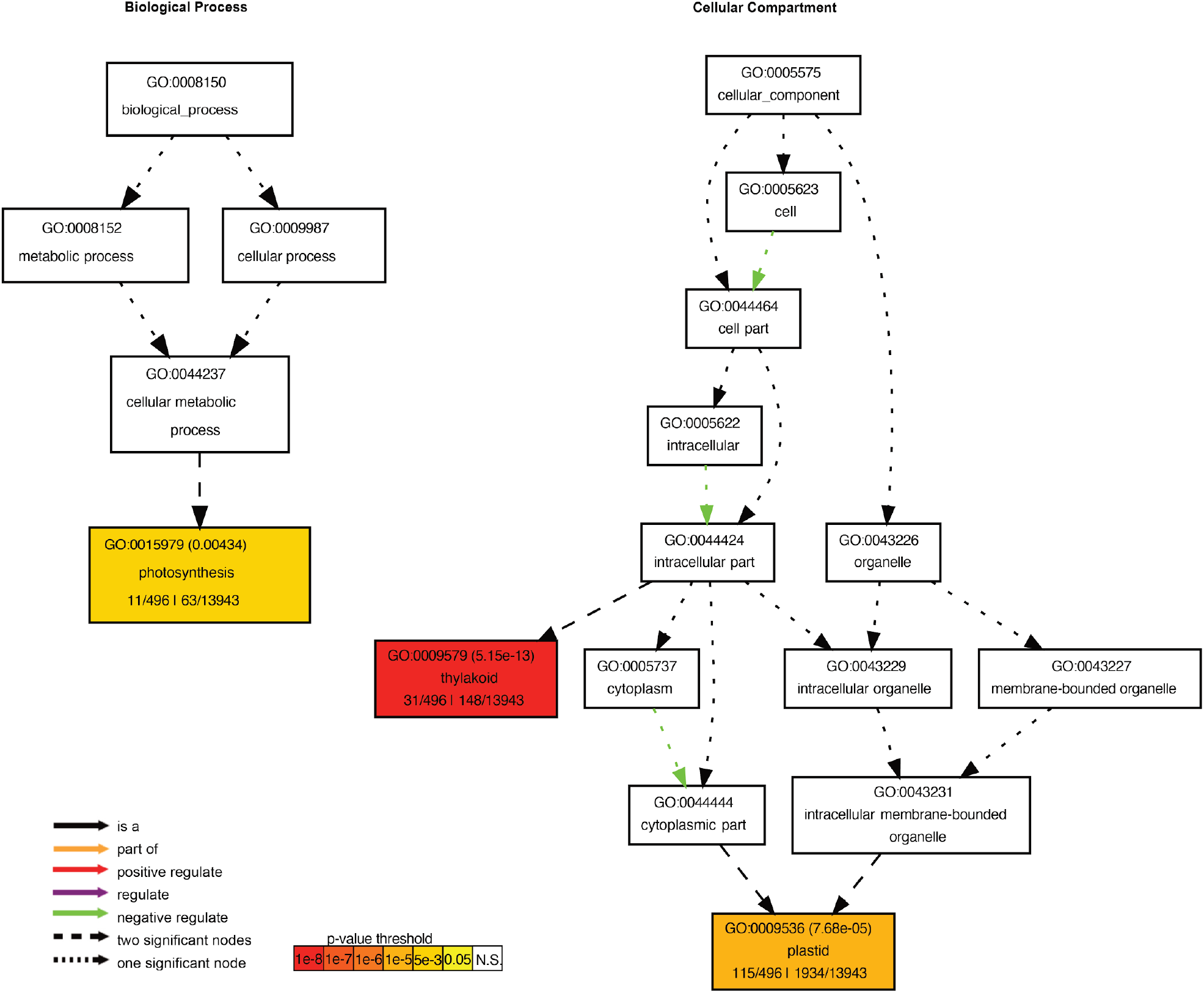
Positive genes are enriched for genes encoding thykaloid-localized proteins that are involved in photosynthetic process. Fisher’s Exact Test was used against the background list of filtered expressed genes in SHB2 dataset, using plant GOslim based on TAIR10 annotation.

**Figure S11.**
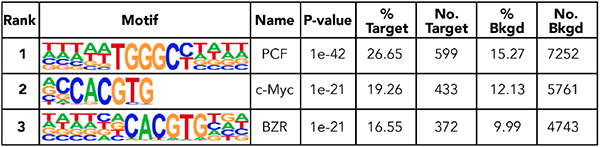
Top 3 motif enrichment results for All-BTH negative genes. All-BTH negative genes are all genes showing negative correlations between dominance of expression with dominance in F1 rosette size when treated with BTH). No.Target: number of genes from All-BTH negative gene list carrying target motif in *cis*, No.Bkgd: number of genes from background gene list carrying target motif in *cis*.

**Figure S12.**
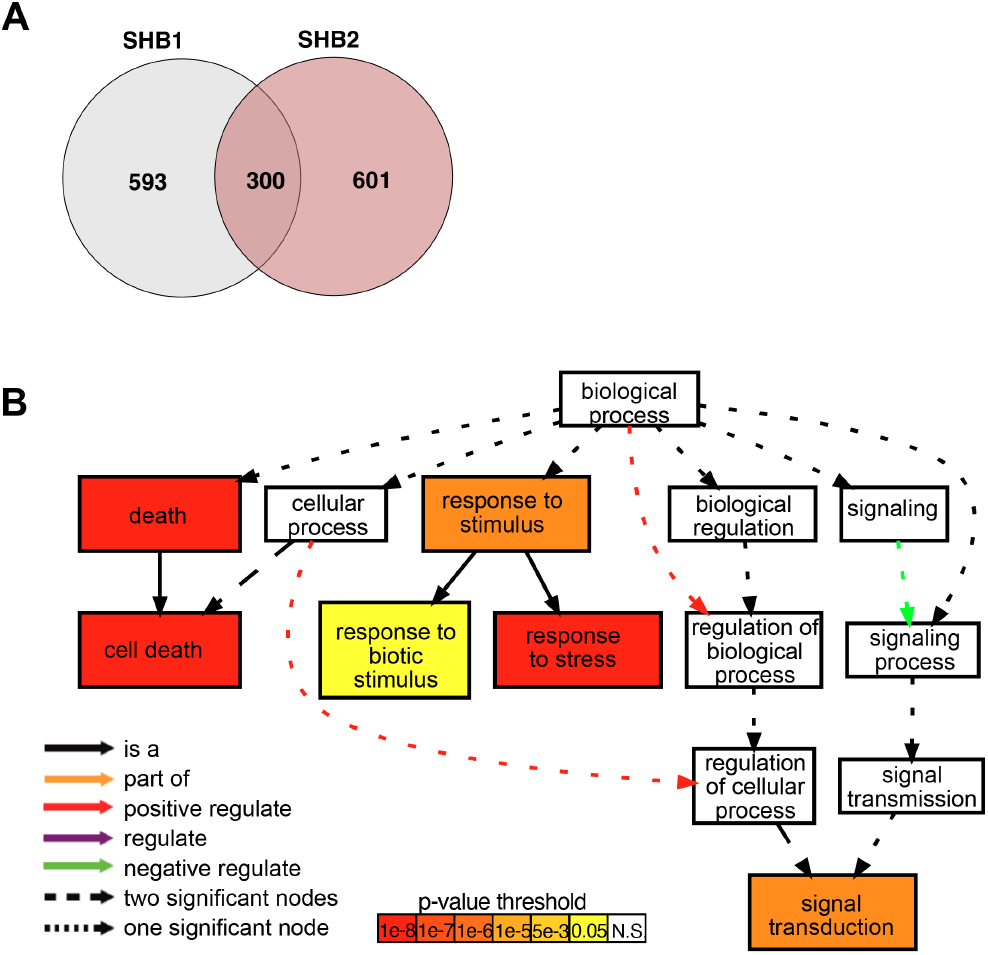
Common additive genes. A. Venn diagram showing the overlapping of additive genes in SHB1 and SHB2 experiments. B. GO-term enrichment diagram of common additive genes.

**Figure S13.**
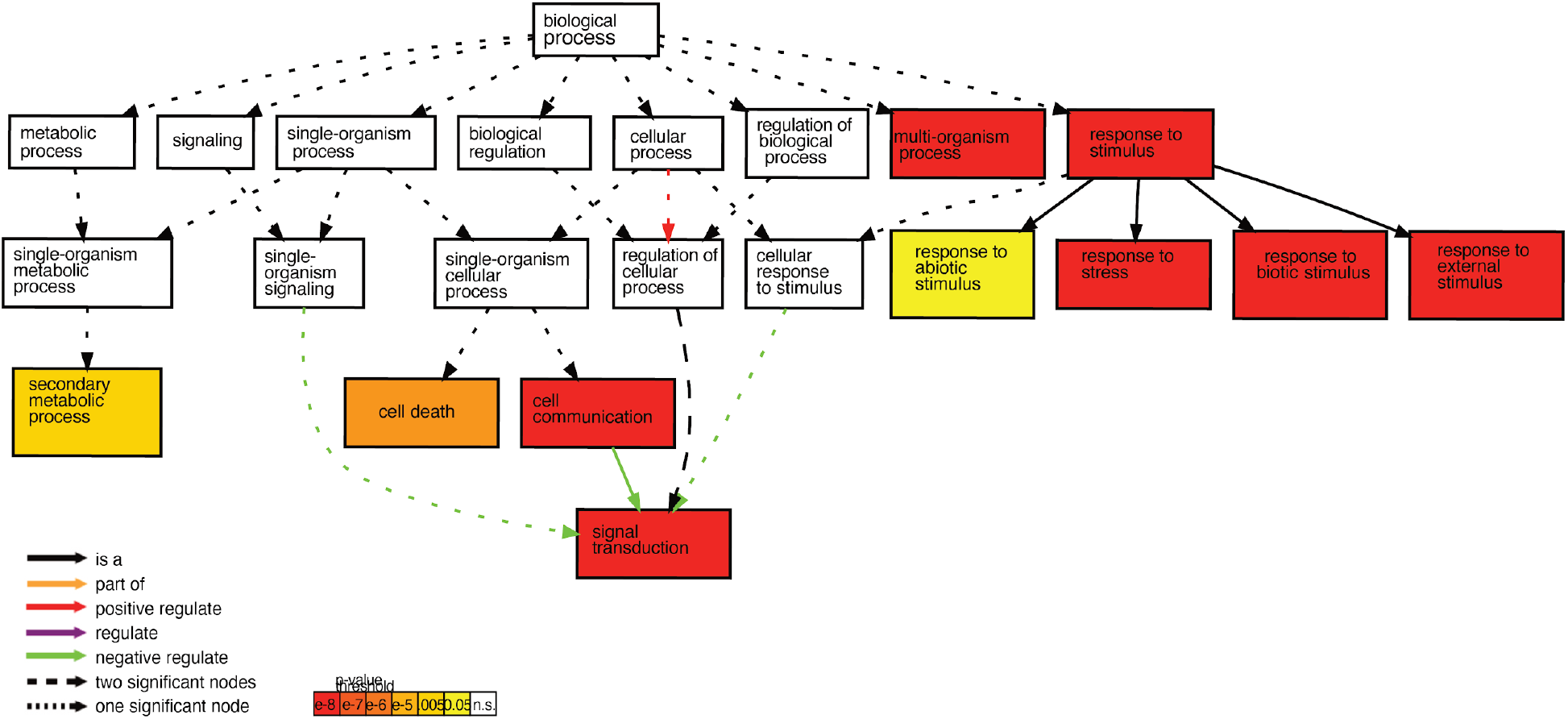
Additive genes in SHB2. This gene set (n=901) is enriched for GO terms of response to stress and (biotic) stimuli, cell death, and secondary metabolic process.

**Fig. S14.**
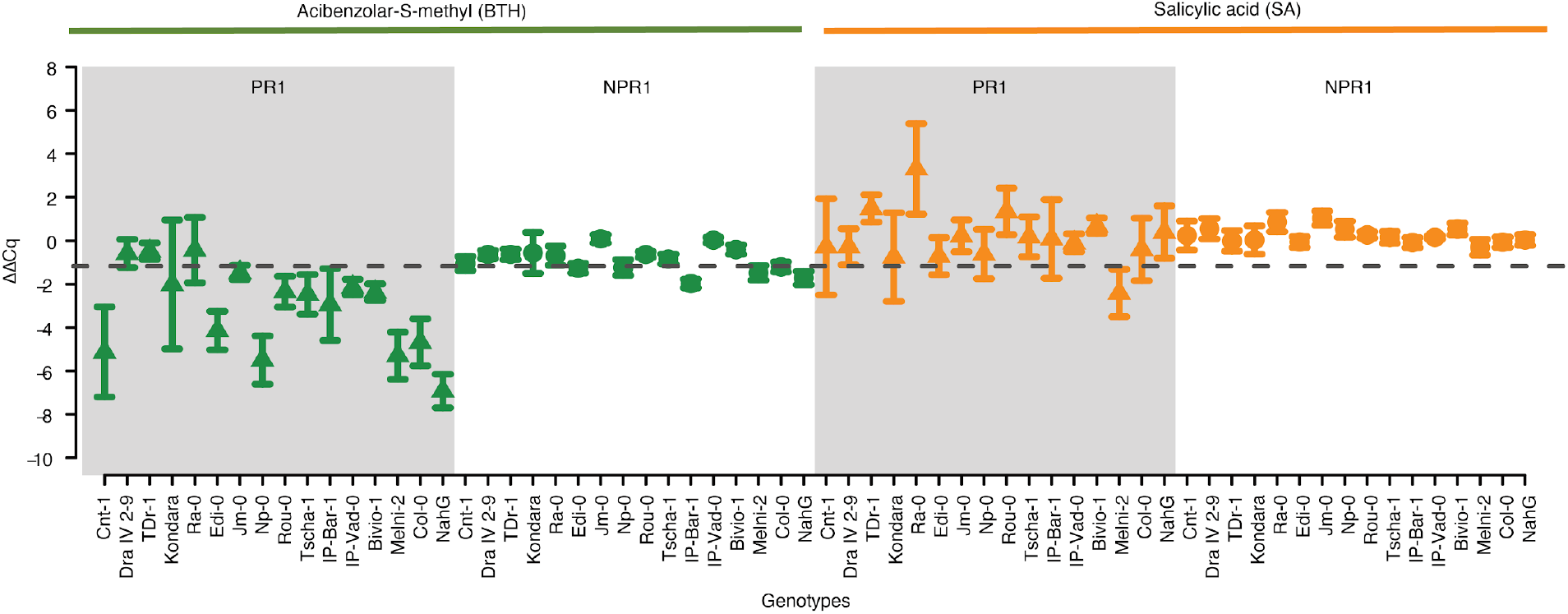
Efficient induction of defense responses in *A. thaliana* accessions with the BTH dosage used. qRT-PCR of two defense marker genes, *PR1* and *NPR1*, in 16 inbred accessions after BTH and SA treatment. Each dot represents the mean ΔΔCq value against housekeeping genes in mock-treated plants, with error bars indicating the standard deviation of the biological replicates.

## References

1001 Genomes Consortium. 2016. 1,135 Genomes Reveal the Global Pattern of Polymorphism in Arabidopsis thaliana. Cell 166:481–491.

Albert FW, Kruglyak L. 2015. The role of regulatory variation in complex traits and disease. Nat. Rev. Genet. 16:197–212.

Andrew Gelman, Yu-Sung Su, Masanao Yajima, Jennifer Hill, Maria Grazia Pittau, Jouni Kerman, Tian Zheng, Vincent Dorie. 2021. Data Analysis Using Regression and Multilevel/Hierarchical Models [R package arm version 1.12-2]. Available from: https://cran.r-project.org/web/packages/arm/index.html

Bates D, Mächler M, Bolker B, Walker S. 2015. Fitting Linear Mixed-Effects Models Using lme4. Journal of Statistical Software, Articles 67:1–48.

Batts GR, Morison JIL, Ellis RH, Hadley P, Wheeler TR. 1997. Effects of CO2 and temperature on growth and yield of crops of winter wheat over four seasons. In: van Ittersum Mk, van de Geijn Sc, editors. Developments in Crop Science. Vol. 25. Elsevier. p. 67–76.

Bell GDM, Kane NC, Rieseberg LH, Adams KL. 2013. RNA-seq analysis of allele-specific expression, hybrid effects, and regulatory divergence in hybrids compared with their parents from natural populations. Genome Biol. Evol. 5:1309–1323.

Berguson WE, McMahon BG, Riemenschneider DE. 2017. Additive and Non-Additive Genetic Variances for Tree Growth in Several Hybrid Poplar Populations and Implications Regarding Breeding Strategy. Silvae Genet. 66:33–39.

Birchler JA, Auger DL, Riddle NC. 2003. In search of the molecular basis of heterosis. Plant Cell 15:2236–2239.

Bomblies K, Lempe J, Epple P, Warthmann N, Lanz C, Dangl JL, Weigel D. 2007. Autoimmune Response as a Mechanism for a Dobzhansky-Muller-Type Incompatibility Syndrome in Plants. PLoS Biol. 5:e236.

Bomblies K, Weigel D. 2007. Hybrid necrosis: autoimmunity as a potential gene-flow barrier in plant species. Nat. Rev. Genet. 8:382–393.

Braendle C, Heyland A, Flatt T. 2011. Integrating mechanistic and evolutionary analysis of life history variation. Mechanisms of life history evolution. The genetics and physiology of life history traits and trade-offs:3–10.

Calleja-Rodriguez A, Chen Z, Suontama M, Pan J, Wu HX. 2021. Genomic Predictions With Nonadditive Effects Improved Estimates of Additive Effects and Predictions of Total Genetic Values in Pinus sylvestris. Front. Plant Sci. 12:666820.

Cambiagno DA, Giudicatti AJ, Arce AL, Gagliardi D, Li L, Yuan W, Lundberg DS, Weigel D, Manavella PA. 2021. HASTY modulates miRNA biogenesis by linking pri-miRNA transcription and processing. Mol. Plant 14:426–439.

Capovilla G, Delhomme N, Collani S, Shutava I, Bezrukov I, Symeonidi E, de Francisco Amorim M, Laubinger S, Schmid M. 2018. PORCUPINE regulates development in response to temperature through alternative splicing. Nat Plants 4:534–539.

Chae E, Bomblies K, Kim S-T, Karelina D, Zaidem M, Ossowski S, Martín-Pizarro C, Laitinen RAE, Rowan BA, Tenenboim H, et al. 2014. Species-wide genetic incompatibility analysis identifies immune genes as hot spots of deleterious epistasis. Cell 159:1341–1351.

Conn SJ, Hocking B, Dayod M, Xu B, Athman A, Henderson S, Aukett L, Conn V, Shearer MK, Fuentes S, et al. 2013. Protocol: optimising hydroponic growth systems for nutritional and physiological analysis of Arabidopsis thaliana and other plants. Plant Methods 9:4.

Crow JF. 2010. On epistasis: why it is unimportant in polygenic directional selection. Philos. Trans. R. Soc. Lond. B Biol. Sci. 365:1241–1244.

Elings A, White JW, Edmeades GO. 1997. Options for breeding for greater maize yields in the tropics. In: van Ittersum Mk, van de Geijn Sc, editors. Developments in Crop Science. Vol. 25. Elsevier. p. 155–168.

Falconer DS, Mackay TFC. 1996. Introduction to quantitative genetics, Longman. Essex, England.

Fujimoto R, Taylor JM, Shirasawa S, Peacock WJ, Dennis ES. 2012. Heterosis of Arabidopsis hybrids between C24 and Col is associated with increased photosynthesis capacity. Proc. Natl. Acad. Sci. U. S. A. 109:7109–7114.

Gao R, Helfant LJ, Wu T, Li Z, Brokaw SE, Stock AM. 2021. A balancing act in transcription regulation by response regulators: titration of transcription factor activity by decoy DNA binding sites. Nucleic Acids Res. 49:11537–11549.

Gonzalez DH. 2015. Plant Transcription Factors: Evolutionary, Structural and Functional Aspects. Academic Press

González R, Butković A, Rivarez MPS, Elena SF. 2020. Natural variation in Arabidopsis thaliana rosette area unveils new genes involved in plant development. Sci. Rep. 10:17600.

Heinz S, Benner C, Spann N, Bertolino E, Lin YC, Laslo P, Cheng JX, Murre C, Singh H, Glass CK. 2010. Simple combinations of lineage-determining transcription factors prime cis-regulatory elements required for macrophage and B cell identities. Mol. Cell 38:576–589.

Hemani G, Knott S, Haley C. 2013. An evolutionary perspective on epistasis and the missing heritability. PLoS Genet. 9:e1003295.

Jeffrey Chen Z. 2013. Genomic and epigenetic insights into the molecular bases of heterosis. Nat. Rev. Genet. 14:471–482.

Julkowska MM, Klei K, Fokkens L, Haring MA, Schranz ME, Testerink C. 2016. Natural variation in rosette size under salt stress conditions corresponds to developmental differences between Arabidopsis accessions and allelic variation in the LRR-KISS gene. J. Exp. Bot. 67:2127–2138.

Korves TM, Schmid KJ, Caicedo AL, Mays C, Stinchcombe JR, Purugganan MD, Schmitt J. 2007. Fitness effects associated with the major flowering time gene FRIGIDA in Arabidopsis thaliana in the field. Am. Nat. 169:E141–E157.

Koşar Z, Erbaş A. 2022. Can the Concentration of a Transcription Factor Affect Gene Expression? Frontiers in Soft Matter [Internet] 2. Available from: https://www.frontiersin.org/articles/10.3389/frsfm.2022.914494

Kremling KAG, Chen S-Y, Su M-H, Lepak NK, Romay MC, Swarts KL, Lu F, Lorant A, Bradbury PJ, Buckler ES. 2018. Dysregulation of expression correlates with rare-allele burden and fitness loss in maize. Nature 555:520–523.

Kumar S, Molloy C, Muñoz P, Daetwyler H, Chagné D, Volz R. 2015. Genome-Enabled Estimates of Additive and Nonadditive Genetic Variances and Prediction of Apple Phenotypes Across Environments. G3 5:2711–2718.

Landry CR, Hartl DL, Ranz JM. 2007. Genome clashes in hybrids: insights from gene expression. Heredity 99:483–493.

Li S. 2015. The Arabidopsis thaliana TCP transcription factors: A broadening horizon beyond development. Plant Signal. Behav. 10:e1044192.

Liu W, He G, Deng XW. 2021. Biological pathway expression complementation contributes to biomass heterosis in Arabidopsis. Proc. Natl. Acad. Sci. U. S. A. [Internet] 118. Available from: http://dx.doi.org/10.1073/pnas.2023278118

Mackay TFC, Stone EA, Ayroles JF. 2009. The genetics of quantitative traits: challenges and prospects. Nat. Rev. Genet. 10:565–577.

Merilä J, Sheldon BC. 1999. Genetic architecture of fitness and nonfitness traits: empirical patterns and development of ideas. Heredity 83 (Pt 2):103–109.

Miller M, Song Q, Shi X, Juenger TE, Chen ZJ. 2015. Natural variation in timing of stress-responsive gene expression predicts heterosis in intraspecific hybrids of Arabidopsis. Nat. Commun. 6:7453.

Misztal I. 1997. Estimation of Variance Components with Large-Scale Dominance Models. J. Dairy Sci. 80:965–974.

Oakley CG, Lundemo S, Ågren J, Schemske DW. 2019. Heterosis is common and inbreeding depression absent in natural populations of Arabidopsis thaliana. J. Evol. Biol. [Internet]. Available from: http://dx.doi.org/10.1111/jeb.13441

Quinlan AR, Hall IM. 2010. BEDTools: a flexible suite of utilities for comparing genomic features. Bioinformatics 26:841–842.

R Core Team. 2021. R: A language and environment for statistical computing. R Foundation for Statistical Computing, Vienna, Austria. Available from: https://www.R-project.org/

Rieseberg LH, Kim S-C, Randell RA, Whitney KD, Gross BL, Lexer C, Clay K. 2007. Hybridization and the colonization of novel habitats by annual sunflowers. Genetica 129:149–165.

Seymour DK, Chae E, Grimm DG, Martín Pizarro C, Habring-Müller A, Vasseur F, Rakitsch B, Borgwardt KM, Koenig D, Weigel D. 2016. Genetic architecture of nonadditive inheritance in Arabidopsis thaliana hybrids. Proc. Natl. Acad. Sci. U. S. A. 113:E7317–E7326.

Su G, Christensen OF, Ostersen T, Henryon M, Lund MS. 2012. Estimating additive and non-additive genetic variances and predicting genetic merits using genome-wide dense single nucleotide polymorphism markers. PLoS One 7:e45293.

Tian T, Liu Y, Yan H, You Q, Yi X, Du Z, Xu W, Su Z. 2017. agriGO v2.0: a GO analysis toolkit for the agricultural community, 2017 update. Nucleic Acids Res. 45:W122–W129.

Varona L, Legarra A, Toro MA, Vitezica ZG. 2018. Non-additive Effects in Genomic Selection. Front. Genet. 9:78.

Vasseur F, Bresson J, Wang G, Schwab R, Weigel D. 2018. Image-based methods for phenotyping growth dynamics and fitness components in Arabidopsis thaliana. Plant Methods 14:63.

Vasseur F, Fouqueau L, de Vienne D, Nidelet T, Violle C, Weigel D. 2019. Nonlinear phenotypic variation uncovers the emergence of heterosis in Arabidopsis thaliana. PLoS Biol. 17:e3000214.

Vogel A, Ebeling A, Gleixner G, Roscher C, Scheu S, Ciobanu M, Koller-France E, Lange M, Lochner A, Meyer ST, et al. 2019. Chapter Seven - A new experimental approach to test why biodiversity effects strengthen as ecosystems age. In: Eisenhauer N, Bohan DA, Dumbrell AJ, editors. Advances in Ecological Research. Vol. 61. Academic Press. p. 221–264.

Wainberg M, Sinnott-Armstrong N, Mancuso N, Barbeira AN, Knowles DA, Golan D, Ermel R, Ruusalepp A, Quertermous T, Hao K, et al. 2019. Opportunities and challenges for transcriptome-wide association studies. Nat. Genet. 51:592–599.

Wellmann R, Bennewitz J. 2012. Bayesian models with dominance effects for genomic evaluation of quantitative traits. Genet. Res. 94:21–37.

Wieters B, Steige KA, He F, Koch EM, Ramos-Onsins SE, Gu H, Guo Y-L, Sunyaev S, de Meaux J. 2021. Polygenic adaptation of rosette growth in Arabidopsis thaliana. PLoS Genet. 17:e1008748.

Xiang T, Christensen OF, Vitezica ZG, Legarra A. 2016. Genomic evaluation by including dominance effects and inbreeding depression for purebred and crossbred performance with an application in pigs. Genet. Sel. Evol. 48:92.

Yang M, Wang X, Ren D, Huang H, Xu M, He G, Deng XW. 2017. Genomic architecture of biomass heterosis in Arabidopsis. Proc. Natl. Acad. Sci. U. S. A. [Internet]. Available from: http://dx.doi.org/10.1073/pnas.1705423114

Zhu Z, Bakshi A, Vinkhuyzen AAE, Hemani G, Lee SH, Nolte IM, van Vliet-Ostaptchouk Jv, Snieder H, LifeLines Cohort Study, Esko T, et al. 2015. Dominance genetic variation contributes little to the missing heritability for human complex traits. Am. J. Hum. Genet. 96:377–385.

Zuk O, Hechter E, Sunyaev SR, Lander ES. 2012. The mystery of missing heritability: Genetic interactions create phantom heritability. Proc. Natl. Acad. Sci. U. S. A. 109:1193–1198.

